# Spatiotemporal dynamics of Mcl-1 abundance and its influence on apoptosis susceptibility

**DOI:** 10.64898/2026.02.06.704006

**Authors:** Fabian Klötzer, Nadine Pollak, Anna Baatz, Batuhan Kisakol, Fiona Ginty, Daniel B. Longley, Jochen H. M. Prehn, Markus Rehm

## Abstract

The Bcl-2 protein family defines cellular competence for mitochondrial outer membrane permeabilization (MOMP) and apoptotic cell death. In proliferating cells, the Bcl-2 family member Mcl-1 accumulates across the cell cycle and confers trans-mitotic resistance to extrinsic apoptosis. We show here that Mcl-1, but not Bcl-xL, additionally undergoes a coordinated redistribution from the cytosol to mitochondria, concomitant with its over-proportional accumulation late in the cell cycle. Live-cell monitoring of Mcl-1 dynamics at single-cell resolution, combined with mathematical modelling, enabled us to quantify that Mcl-1 redistribution substantially contributes to elevating MOMP thresholds. Furthermore, we found that Mcl-1 accumulation and redistribution act concomitantly but independently to increase MOMP thresholds as cells approach mitosis and this elevated resistance is reset in daughter cells after division. Notably, heterogeneities in Mcl-1 abundance and subcellular distribution are pronounced even among isogenic cells within the same cell-cycle phase, and thus contribute to substantial cell-to-cell variability in MOMP susceptibility. Analysis of colorectal cancer tissue samples showed that variability in Mcl-1 expression and distribution is likewise prominent between cells in patient tumors and were predicted to drive intra-tumour heterogeneity in responses to treatments that induce MOMP. Overall, we demonstrate how changes in Mcl-1 amounts and localisation integrate with cell-cycle progression to modulate apoptotic susceptibility, thereby shaping cell-fate outcomes and contributing to cell-to-cell heterogeneities in death decision making.

## Introduction

Apoptosis is a highly regulated form of programmed cell death, which is required during development, for elimination of damaged or infected cells and for maintaining tissue homeostasis (Nagata & Tanaka, 2017). Consequently, dysregulated apoptosis can contribute to degenerative disorders and proliferative diseases, such as cancer (Hanahan, 2022; Vitale et al., 2023). Many different stressors and types of cellular damage can initiate apoptosis signalling through the extrinsic or intrinsic apoptosis pathways, both of which converge at the mitochondria, where sufficient apoptotic signals lead to mitochondrial outer membrane permeabilization (MOMP). MOMP represents a central checkpoint that triggers the execution phase of apoptosis and is both orchestrated and tightly regulated through the interactions of members of the Bcl-2 protein family (Czabotar & Garcia-Saez, 2023; Singh et al., 2019). The pro-apoptotic members Bax and Bak can oligomerize to form pores in the outer mitochondrial membrane, whereas anti-apoptotic inhibitors such as Bcl-2, Bcl-xL and Mcl-1 inhibit Bax and Bak. Members of the group of pro-apoptotic BH3-only proteins either antagonize the anti-apoptotic Bcl-2 family proteins or additionally directly activate Bax and Bak (Czabotar & Garcia-Saez, 2023; Singh et al., 2019). Elevated expression of anti-apoptotic Bcl-2 family members consequently promotes apoptosis resistance by neutralizing BH3-only proteins and keeping Bax and Bak in check, and overexpression or gene amplification of Bcl-2, Bcl-xL and Mcl-1 is frequently found in various cancers (Beroukhim et al., 2010; Tsujimoto et al., 1984). Despite their overlapping protective functions, these proteins differ markedly in their regulation and stability. Bcl-2 and Bcl-xL are relatively stable and provide sustained apoptosis protection (Goodall et al., 2016). In contrast, Mcl-1 is turned over far more rapidly (half-life = 20 – 40 min, (Nijhawan et al., 2003; Schwickart et al., 2010)), allowing its protein levels to be rapidly adjusted by changes in transcription, translation, and proteasomal degradation (Senichkin et al., 2020). Mcl-1 is predominantly localized to the MOM, but Mcl-1 is also reported to be found in the cytoplasm, at the endoplasmic reticulum, and in the nucleus (Fu et al., 2022; Kale et al., 2018; Nakajima et al., 2014). At the MOM, Mcl-1 exerts its canonical anti-apoptotic function by sequestering pro-apoptotic pore formers, but also appears to support mitochondrial bioenergetics and cellular metabolic fitness (Brinkmann et al., 2025; Wright et al., 2024). In contrast to Bcl-2 and Bcl-xL, Mcl-1 expression is additionally regulated by cell cycle progression (Harley et al., 2010; Pollak et al., 2021) with elevated Mcl-1 amounts conferring transient, trans-mitotic resistance to extrinsic apoptosis (Pollak et al., 2021).

Here, we combined quantitative, high-resolution imaging at single cell and subcellular scales together with mathematical modelling to elucidate how Mcl-1 expression dynamics and subcellular distribution define mitochondrial apoptosis thresholds. Furthermore, we show how cell-to-cell differences in apoptosis susceptibility emerge from natural intercellular heterogeneities in Mcl-1 expression and distribution, and demonstrate that both factors independently contribute to apoptosis resistance.

## Results

### Mcl-1 accumulates with progression in cell cycle and redistributes to mitochondria

Previous biochemical analyses in cell populations and single-cell microscopic analyses showed that Mcl-1 levels are regulated across the cell cycle (Harley et al., 2010; Pollak et al., 2021). Moreover, maximum intensity projection-imaging indicated that Mcl-1 might redistribute towards the mitochondria during cell cycle progression, although these measurements lacked authentic spatial resolution (Pollak et al., 2021). Here, we therefore set out to quantitatively determine genuine relative changes in Mcl-1 amounts and localisation at single cell resolution within and across major cellular compartments, including the nucleus, the cytosol and mitochondria. NCI-H460 cells expressing geminin-GFP as a cell cycle indicator were labelled with MitoTracker red and DAPI as segmentation markers for mitochondria and nuclei, respectively, and stained for Mcl-1 (**Figure 1A**). Single layer segmentation allowed us to separate nuclear, cytosolic and mitochondrial regions (**Figure 1B**, see also methods section). Mcl-1 concentrations were highest in mitochondrial regions, corresponding to results for Bcl-xL, a related Bcl-2 family member (**Figure 1C**). The presence of non-mitochondrial pools of Mcl-1 was confirmed also by its signals substantially exceeding those obtained by using secondary antibody-only staining, as well as by biochemical fractionation (**Figure S1A-C**). Separating and comparing cells from early and late cell cycle stages based on absent or very high geminin reporter signals (**Figure S1D**) showed that Mcl-1 concentrations in all compartments increased from early cell cycle (G_1_) to late cell cycle stages (late S/G_2_), with the highest concentration increases observed at the mitochondria (**Figure 1D, E**). In contrast, subcellular Bcl-xL concentrations did not increase in any of the compartments (**Figure 1F**). To compare subcellular protein distributions of Mcl-1, we calculated the ratios between cytoplasmic and mitochondrial mean intensities for each individual cell. While ratios for Bcl-xL remained constant, Mcl-1 very prominently redistributed towards mitochondria (**Figure 1G**). While the size of nuclear, cytosolic and mitochondrial areas all increased across the cell cycle (**Figure S1E**), Mcl-1 concentrations and distributions did not correlate with and therefore could not be explained by these changes (**Figure S1F-H**). Taken together, these analyses thus demonstrate that cellular Mcl-1 concentrations increase with cell cycle progression and additionally that Mcl-1 disproportionally accumulates at the mitochondria (**Figure 1H**).

**Figure 1:**
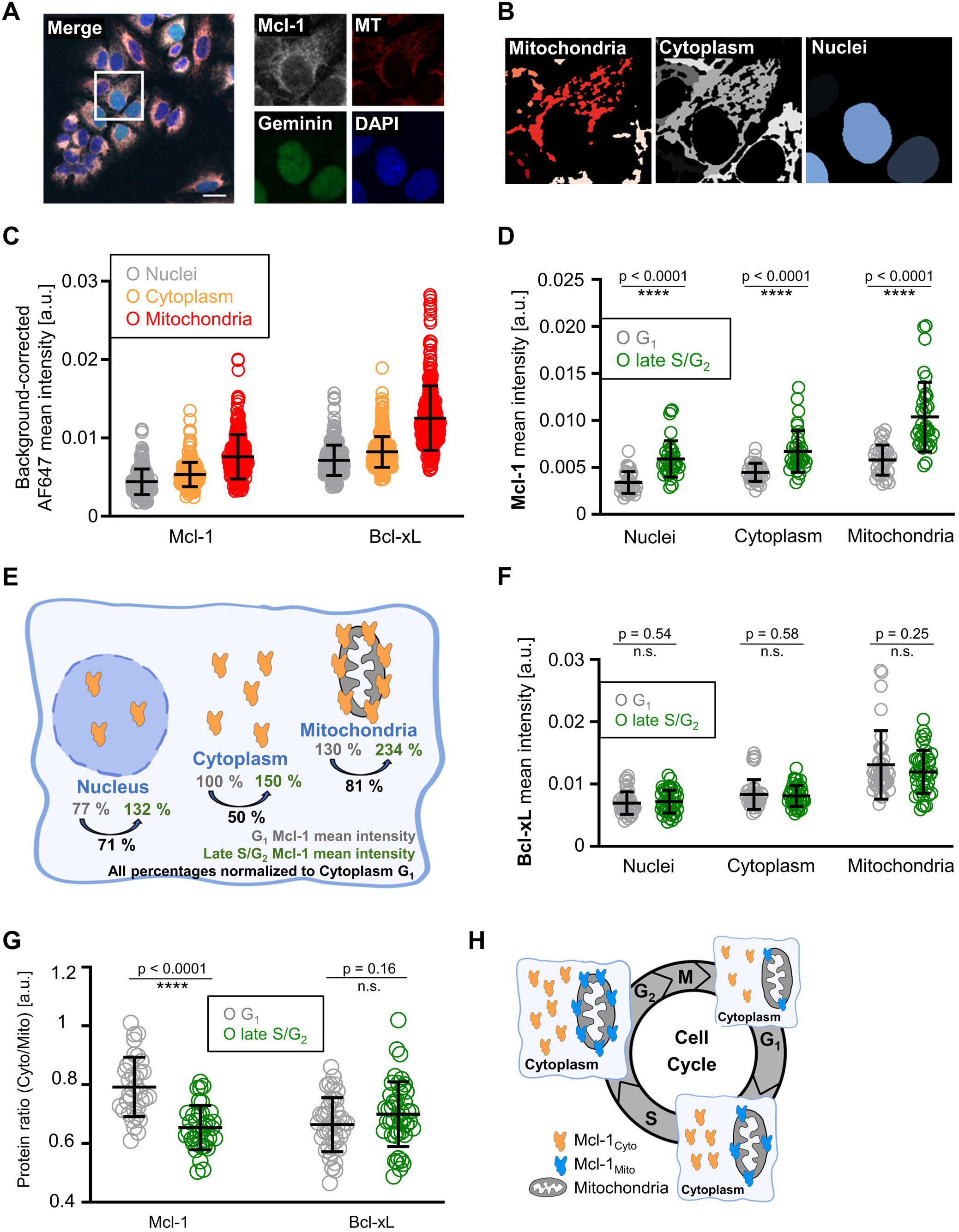
Mcl-1 accumulates with progression in cell cycle and redistributes to mitochondria. A) NCI-H460/geminin cells were incubated with MitoTracker (red), fixed and stained for Mcl-1 (white) and DNA (blue). Scale bar = 20 µm. B) Segmentation of mitochondrial, cytoplasmic and nuclear area for a representative cell. C) Mean Mcl-1 or Bcl-xL intensities were background-corrected and measured within the segmented compartments. n = 243 cells for Mcl-1 and n = 347 cells for Bcl-xL. D) Comparison of subcellular Mcl-1 concentrations between cells from early and late in the cell cycle, based on geminin expression intensities. n = 35 cells per group. E) Relative increases in Mcl-1 concentrations in the respective compartments with cell cycle progression. F) Comparison of subcellular Bcl-xL concentrations between cells from early and late in the cell cycle, based on geminin expression intensities. n = 44 cells per group. G) Protein distributions of Mcl-1 and Bcl-xL early and late in the cell cycle, displayed as their cytoplasmic to mitochondrial ratios. n = 35 cells per group for Mcl-1 and 44 cells per group for Bcl-xL. H) Scheme depicting the simultaneous increase in Mcl-1 expression and its redistribution to the mitochondria with progression in cell cycle. (C,D,F,G) show one representative experiment from 3 independent biological repeats. Each dot represents a single cell. Error bars represent mean ± standard deviation. *p*-values from an unpaired t-test with Welch’s correction are depicted.

### Mcl-1 redistribution occurs concomitantly with, but independently of, Mcl-1 accumulation

To study whether or not increases in Mcl-1 across the cell cycle and its redistribution towards mitochondria are temporally and mechanistically coupled, we applied a combinatorial cell staining and pseudotime analysis. To infer the dynamics of changes in Mcl-1 expression and redistribution, we combined geminin-GFP signals and DAPI intensities, the latter indicating DNA content. As snapshot data, both provide information on the cell cycle position of individual cells from S-phase into G_2_ and on towards entry into mitosis. These positions were then linked to cellular Mcl-1 staining data. Pooling and normalizing independent experiments (**Figure 2A**) provided us with information from a total of 4500 individual cells. Mcl-1 amounts and Mcl-1 redistribution to the mitochondria concomitantly increased along both the geminin-GFP and DAPI dimensions (**Figure 2B,C**). Due to the short duration of mitosis, M phase cells were quantitatively underrepresented in this analysis. We therefore separately collected imaging data including cells in M phase and newly divided G_1_ sibling cells (**Figure 2D**). Cells in M phase expressed Mcl-1 at significantly higher concentrations than in S/G_2_ phases (**Figure 2E**). Cells right after or during division initially retained the high Mcl-1 concentration observed in M phase (**Figure 2E**). Cells also maintained the mitochondrial accumulation of Mcl-1 during mitosis and in newly divided sibling cells (**Figure 2F**). This suggests that daughter cells initially inherit both increased expression and mitochondrial accumulation of Mcl-1 from the mother cells, before Mcl-1 amounts and distributions revert back to values found in typical G_1_ cells. Mcl-1 expression and redistribution are therefore also regulated concurrently during and across mitosis.

**Figure 2:**
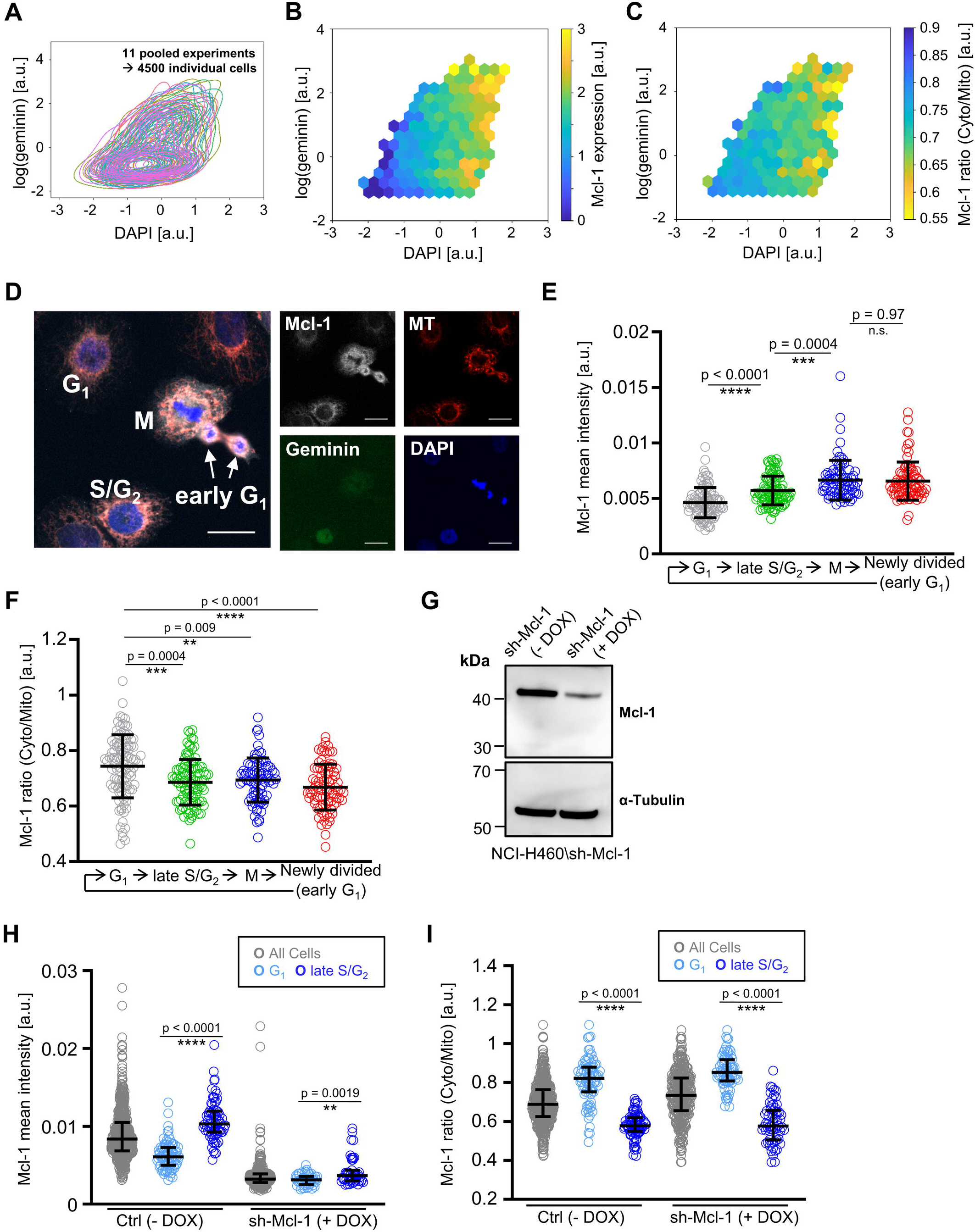
Mcl-1 redistribution occurs concomitantly with but independently of Mcl-1 accumulation. A) Data from 11 individual experiments (NCI-H460/geminin cells were incubated with MitoTracker, fixed and stained for Mcl-1 and DNA) were pooled. The pooled, normalized distributions are shown. B) Hexbin plot of geminin vs. DAPI intensities as a proxy for cell cycle progression. The colour code indicates the Mcl-1 expression levels. C) Hexbin plotting as in (B), with the colour indicating the Mcl-1 distribution between cytoplasm and mitochondria. D) NCI-H460/geminin cells were incubated with MitoTracker, fixed, and stained for Mcl-1 and DAPI, with a focus on M phase cells and newly divided sibling cells (very early G_1_). Scale bar = 20 µm. E) Mcl-1 mean intensities in different cell cycle stages were quantified. n > 80 cells per group. F) Mcl-1 distribution ratios in different cell cycle stages were quantified. G) Immunoblot showing the DOX-inducible downregulation of Mcl-1 expression in NCI-H460/sh-Mcl-1 cells. H) Mcl-1 mean intensities from untreated cells and cells with DOX-induced Mcl-1 downregulation are depicted and compared between G_1_ and late S/G_2_ phases. I) Mcl-1 distribution ratios from untreated cells and cells with DOX-induced Mcl-1 downregulation are depicted and compared between G_1_ and late S/G_2_ phases. (E, F) show one representative experiment from two independent biological repeats. Each dot represents a single cell. Error bars represent mean ± standard deviation. *p*-values from a One-Way ANOVA with Dunnett’s T3 multiple comparisons are depicted. (H, I) show one representative experiment from three independent biological repeats. Each dot represents a single cell. Error bars represent median ± interquartile range. *p*-values from Mann-Whitney tests are depicted.

We next assessed if accumulation and redistribution depended on each other. NCI-H460 cells with inducible Mcl-1 depletion (**Fig. 2G**) failed to accumulate notable amounts of Mcl-1 from G_1_ to S/G_2_ phases (**Fig.2H**). However, the low amounts of remaining Mcl-1 still redistributed towards the mitochondria (**Figure 2I**). This finding excludes that the high accumulation of Mcl-1 in late cell cycle phases alone is already sufficient to drive its over-proportional accumulation at the mitochondria.

Taken together, we therefore conclude that both Mcl-1 expression levels and its subcellular distribution are regulated in parallel with cell cycle progression, yet independently of each other.

### Mcl-1 is rapidly exchanged between mitochondria and cytoplasm

The detection of Mcl-1 amounts at the mitochondria, but also in the cytoplasm and nucleus, and the evidence for its spatiotemporal redistribution raise the question on how mobile Mcl-1 is between these compartments.

To study exchange kinetics between mitochondria and cytoplasm, we fluorescently tagged endogenous Mcl-1 with mScarlet at the N-terminus via CRISPR-Cas9 knock-in in NCI-H460 cells (H460-S-Mcl-1). H460-S-Mcl-1 cells expressed mScarlet-Mcl-1 at the expected molecular weight (**Figure 3A,B**). Sequencing and PCR amplification further confirmed the correct insertion of mScarlet DNA into the Mcl-1 locus (**Figure S2A**), and mScarlet-Mcl-1 fluorescence dropped when targeting Mcl-1 expression by siRNA (**Figure S2B**). Furthermore, mScarlet-Mcl-1 expression increased and mScarlet-Mcl-1 redistributed towards the mitochondria along progression in the cell cycle (**Figure 3C,D, Figure S2C**). mScarlet-Mcl-1 maintained its antiapoptotic function, as shown by comparable time-to-death profiles between NCI-H460 WT and H460-S-Mcl-1 cells upon Fc-scTRAIL treatment, and by eliminating the previously described transmitotic resistance to TRAIL treatment (Pollak et al., 2021) by pharmacological inhibition of mScarlet-Mcl-1 (**Figure S2D,E**). Taken together, these findings demonstrate that the mScarlet-Mcl1 fusion protein authentically reflects the behaviour and anti-apoptotic potency of the intrinsic untagged protein.

**Figure 3:**
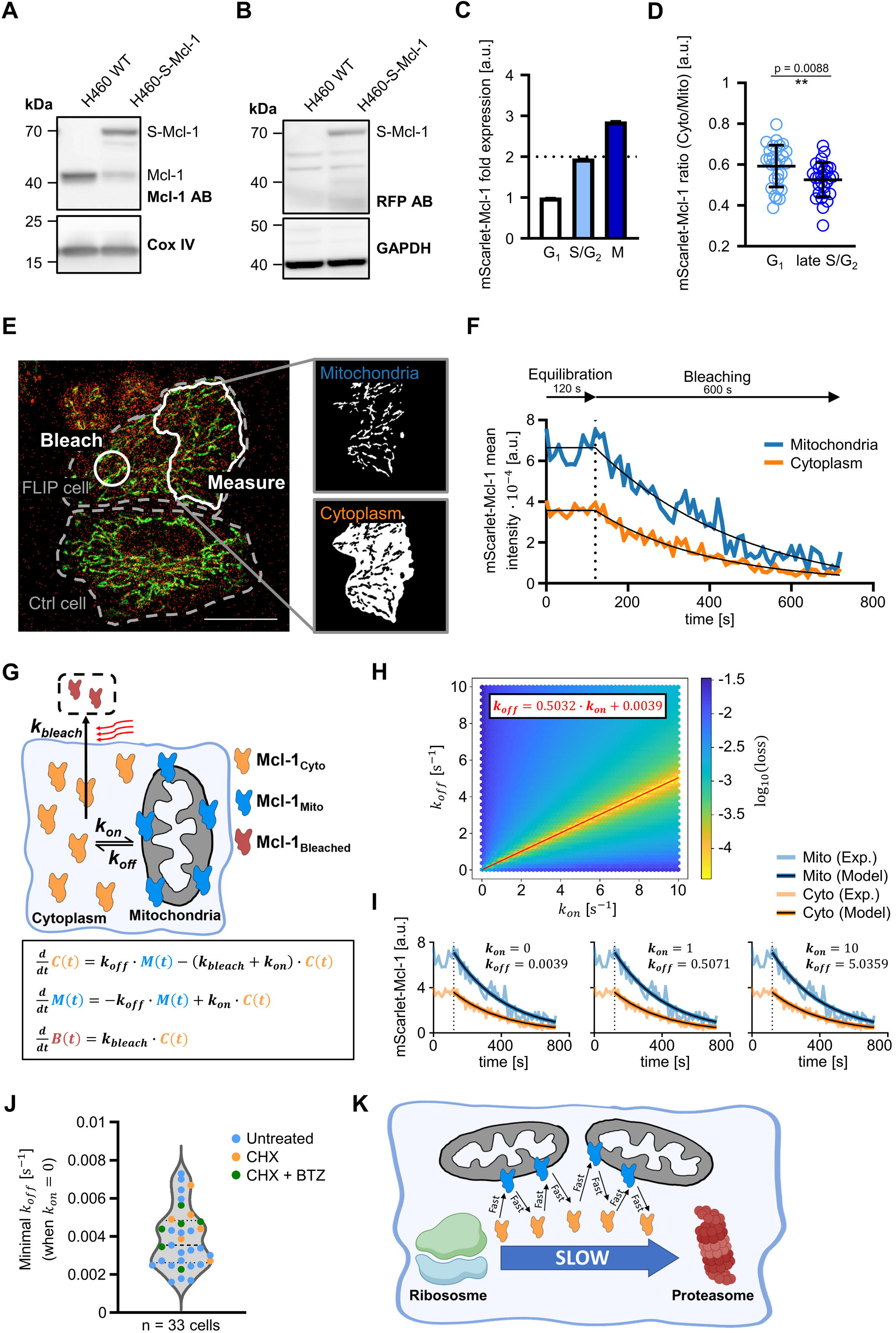
Mcl-1 is rapidly exchanged between mitochondria and cytoplasm. A) Immunoblots showing WT Mcl-1 (41 kDa) or mScarlet-Mcl-1 (70 kDa) expression. B) Immunoblot of mScarlet-Mcl-1 (70 kDa) by RFP-directed antibody detection. C) Fluorescence of mScarlet-Mcl-1 was measured for the different cell cycle stages, separated as shown in Figure S2C. Intensities were normalized to G_1_ and displayed as fold expression relative to G_1_. One representative experiment from two independent repeats is shown. D) H460-S-Mcl-1 cells were imaged for mScarlet-Mcl-1 distribution ratios. Cell cycle stages were discriminated by DNA content using NucBlue dye. n = 29 cells per group, one representative experiment from two independent repeats is shown. E) Exemplary overview of fluorescence loss in photobleaching (FLIP) imaging. H460-S-Mcl1 cells were stained with MitoTracker green. mScarlet-Mcl-1 (red) was bleached in the bleaching area and measured in the segmented measurement areas. Scale bar = 20 µm. F) Fluorescence decay of mScarlet-Mcl-1 fluorescence intensity in the measurement areas. The black lines depict plateaus followed by exponential decays fitted to the experimental data. G) Scheme and underlying ODEs of the mathematical model that describes the FLIP experiments. H) Graph that shows the loss value (difference of the model to the experimental data) depending on the respective combination of 𝑘_𝑜𝑛_ and 𝑘_𝑜𝑓𝑓_ in the FLIP model. The red line depicts a linear function fitted through the optimal combinations, with the equation displayed in the graph. I) Examples of combinations of 𝑘_𝑜𝑛_ and 𝑘_𝑜𝑓𝑓_ along the linear optimum. The model results (black) replicate the experimental data. J) The lowest possible 𝑘_𝑜𝑓𝑓_ values from 33 FLIP analyses. Cells were either left untreated, subjected to 25 µg/ml cycloheximide (CHX) alone, or to a combination of 25 µg/ml cycloheximide and 65 µM bortezomib (BTZ). K) Scheme visualizing the different timescales of turnover and mitochondrial-cytoplasmic exchange.

Next, we measured the exchange kinetics of mScarlet-Mcl-1 (mitochondrial binding and unbinding) by Fluorescence Loss In Photobleaching (FLIP) in single cells (**Figure 3E**). Neighbouring cells served as control cells to correct signals for photobleaching in adjacent regions by light scatter (**Figure S2F-H**). Corrected FLIP measurement data (**Figure S2I**) then allowed us to plot fluorescence decay in the cytoplasm and the mitochondria. Cytoplasmic mScarlet-Mcl-1 intensities declined fast, as expected from free diffusion between the bleaching and the measurement areas (**Figure 3F**). Surprisingly, also mitochondrial mScarlet-Mcl-1 intensity decayed swiftly, suggesting a rather high mobility also for the pool of Mcl-1 associated with mitochondria (**Figure 3F**). To quantify the exchange kinetics, we defined a mathematical model based on ordinary differential equations to describe the FLIP results by estimates of the underlying kinetic parameters (**Figure 3G**). As anticipated, the bleaching rate 𝑘_𝑏𝑙𝑒𝑎𝑐ℎ_ was uniquely identifiable from the experimental data, as it was independent of the other parameters (**Figure S2J**). In contrast, the exchange rates 𝑘_𝑜𝑛_ and 𝑘_𝑜𝑓𝑓_ depended linearly on each other (**Figure 3H**), so that any combination of 𝑘_𝑜𝑛_ and 𝑘_𝑜𝑓𝑓_ along the linear optimum satisfied the experimental data (**Figure 3I**). For simplicity, we set 𝑘_𝑜𝑛_ to zero to determine the minimal 𝑘_𝑜𝑓𝑓_ as an explicit number for the model. Importantly, inhibiting protein translation or degradation by cycloheximide or bortezomib did not affect the range of minimal 𝑘_𝑜𝑓𝑓_ values determined from individual cells in these experiments (**Figure 3J**). Overall, these results therefore demonstrate that Mcl-1 exchanges between the cytoplasmic and mitochondrial pools, and that these exchanges proceed at rates faster than those for protein production or degradation (**Figure 3K**).

To determine if this exchange encompasses the entire Mcl-1 pool, we examined mitochondrial Mcl-1 retention after selective plasma membrane permeabilization with digitonin (Niklas et al., 2011). After permeabilization, large amounts of Mcl-1 diffused out of the cell, yet less than would be expected from an entirely soluble cytosolic protein (compare to GAPDH) (**Figure S3A**). In comparison to Mcl-1, TOM20 signals were far better retained, as expected from a membrane integrated protein (**Figure S3A**). Longer incubation times with digitonin only marginally increased Mcl-1 loss, showing that the remaining Mcl-1 pool is stably retained inside of cells (**Figure S3B**). Observing the loss of Scarlet-tagged Mcl-1 upon digitonin addition in real time provided similar results. A substantial pool of Mcl-1 was lost immediately upon digitonin-based permeabilization, and ratiometric analysis showed that the remaining Mcl-1 was predominantly associated with mitochondria (**Figure S3C,D**). These data suggest, that a fraction of Mcl-1 is quickly washed out of the cell after permeabilization, and that a pool of membrane-associated or integrated Mcl-1 remains stably retained even after longer digitonin incubation. When purifying mitochondria for carbonate extraction experiments, Mcl-1 was not found in supernatants of isolated mitochondria, further confirming that any remaining Mcl-1 is stably associated with or integrated into the outer mitochondrial membrane (**Figure S3E,F**). Carbonate extraction upon elevated pH more easily transferred subpools of Mcl-1 into the supernatant than membrane integrated TOM20, and substantial amounts of Mcl-1 remained tightly associated with the mitochondrial fraction (**Figure S3F**).

Taken together, these results therefore show that Mcl-1 exists in two pools, one of which can rapidly exchange between mitochondria (or mitochondria-associated regions) and the cytosol, the other of which being tightly membrane associated or integrated at the mitochondria.

### Quantitative estimates of altered MOMP thresholds indicate that peak resistance requires both redistribution and accumulation of Mcl-1

Since Mcl-1 accumulation and redistribution occur in parallel during cell cycle progression, we used deterministic mathematical modelling to quantify their relative contribution to altering MOMP susceptibility throughout the cell cycle. We developed a first model to describe spatiotemporal Mcl-1 regulation (**Figure 4A**), to then couple this with a separate model component that allowed to quantify MOMP thresholds. Progression through the cell cycle was integrated by inferring the duration of the individual phases, the proportion of cells in the respective phases and the overall duration of the cell cycle in H460 cells (**Figure S4A**). Furthermore, cellular volume changes with progression in the cell cycle were taken into account (**Figure S4B**). The resulting model could then be used to simulate cell cycle-dependent accumulation of Mcl-1 and its subcellular redistribution (**Figure 4B,C**). To quantitatively verify the accuracy of the model, we simulated a population of 2000 unsynchronized cells (**Figure S4C**) from which population snapshots could directly be compared to experiments. The simulation results for cell cycle-dependent Mcl-1 redistribution and accumulation excellently corresponded to experimental data (**Figure 4D,E**). We then used the model to mathematically uncouple the otherwise biologically concomitant accumulation of Mcl-1 and its redistribution to mitochondria, so that their respective contributions to MOMP thresholds could be estimated independently of one another. In contrast to the full model that reflected both Mcl-1 redistribution and accumulation as observed experimentally (**Figure 4F**), the “Only Redistribution” model variant kept Mcl-1 amounts constant to isolate the sole effect of Mcl-1 redistribution (**Figure 4G**). Conversely, the “Only Accumulation” model variant kept the affinity of Mcl-1 towards the mitochondria constant, thereby separating the accumulation of Mcl-1 throughout the cell cycle from effects arising from altered subcellular redistribution (**Figure 4H**).

**Figure 4:**
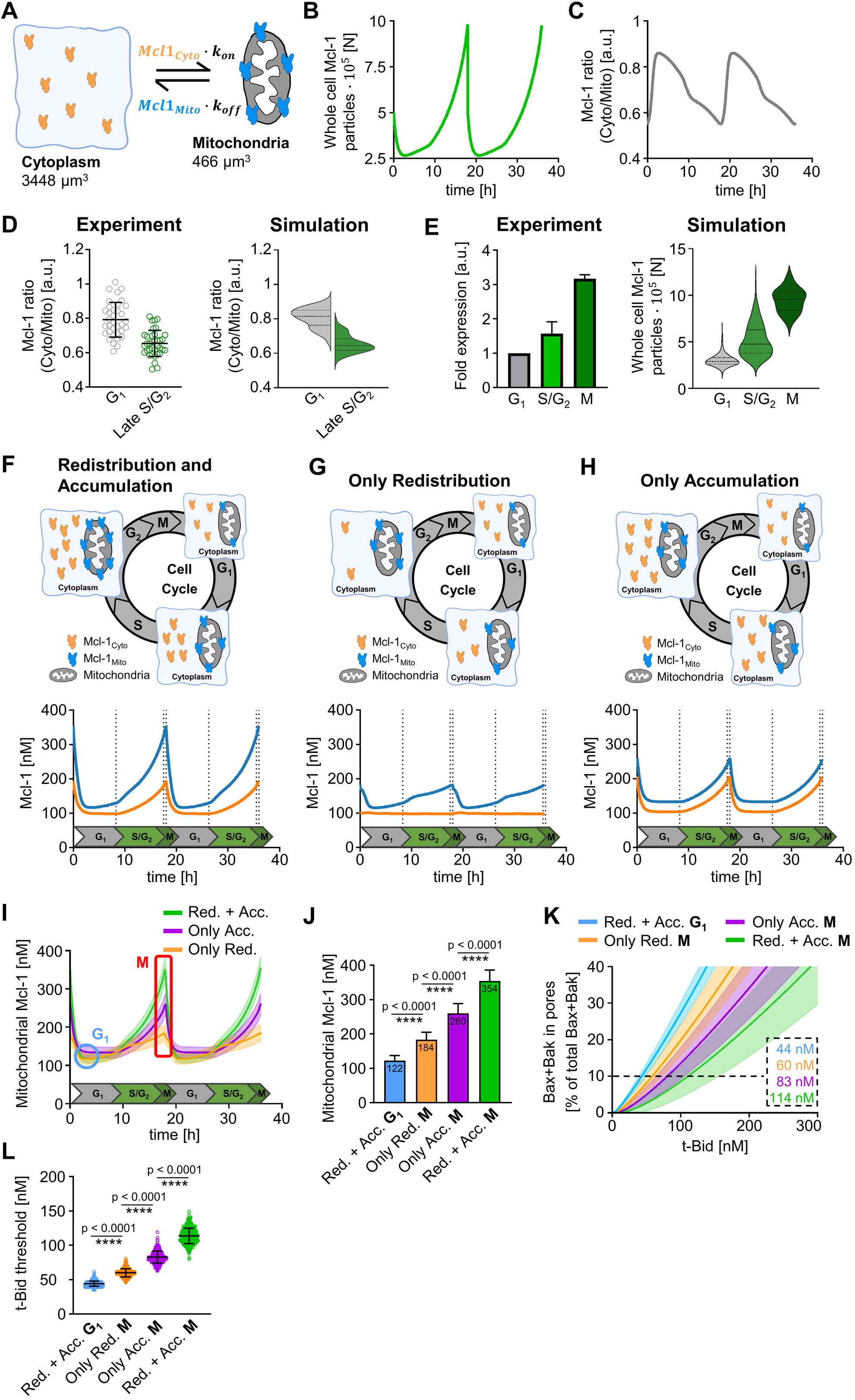
Quantitative estimates of altered MOMP thresholds indicate that peak resistance requires both redistribution and accumulation of Mcl-1. A) Scheme depicting the mitochondrial and cytoplasmic compartments as the foundation of the simulation approach. Mcl-1 exchanges between the two compartments following mass-action kinetics. B) Simulation of Mcl-1 amounts over time. C) Simulation of subcellular Mcl-1 distribution over time. D) Comparison of experimentally measured Mcl-1 distribution ratios and simulated Mcl-1 distribution ratios between early and late cell cycle phases. E) Comparison of experimentally measured Mcl-1 levels and simulated Mcl-1 expression levels in different cell cycle stages. F) Top: Scheme depicting the regulation of Mcl-1 across the cell cycle. Bottom: Mitochondrial (blue) and cytoplasmic (orange) Mcl-1 concentrations simulated by the ODE model over two consecutive cell cycles. G) Top: Scheme depicting the “Only Redistribution” model of Mcl-1. Bottom: Mitochondrial (blue) and cytoplasmic (orange) Mcl-1 concentrations simulated by the ODE model over two consecutive cell cycles. H) Top: Scheme depicting the “Only Accumulation” model of Mcl-1. Bottom: Mitochondrial (blue) and cytoplasmic (orange) Mcl-1 concentrations simulated by the ODE model over two consecutive cell cycles. I) Mitochondrial Mcl-1 concentrations from the models shown in (F-H) are summarized over two cell cycles. Early G_1_ (blue) and M phase (red) are highlighted. For each model, 500 cells reflecting experimentally observed Mcl-1 heterogeneity were simulated. J) Mitochondrial Mcl-1 concentrations in G_1_ were compared to Mcl-1 in M phase in the different models. Error bars represent mean ± standard deviation from the 500 single cells simulated in (I). *p*-values from a One-Way ANOVA with Dunnett’s T3 multiple comparisons post hoc test are depicted. K) Mitochondrial Mcl-1 concentrations from (J) were used as input for an apoptosis susceptibility model. It is shown how much t-Bid in each scenario is necessary to push simulated cells above a threshold of 10% of Bax and Bak in pores. L) The 10% t-Bid threshold values of each scenario are compared using One Way ANOVA with Dunnett’s T3 multiple comparisons. Error bars represent mean ± standard deviation from the 500 single cells simulated in (I).

Mitochondrial Mcl-1 concentrations were very similar between all variants during G_1_, but differed substantially towards the end of the cell cycle (**Figure 4I**). Both redistribution and the accumulation of Mcl-1 significantly contributed to elevating mitochondrial Mcl-1 concentrations, with both acting together to achieve peak mitochondrial Mcl-1 concentrations in M phase (**Figure 4J**). The mitochondrial Mcl-1 concentrations from the model variants then served as input for a simplified, experimentally validated mathematical model capable of estimating MOMP susceptibility (Hantusch et al., 2018). In this model, t-Bid served as a representative and tuneable pro-apoptotic input signal, whereas MOMP susceptibility was assessed by the amount of Bax and Bak in pores (interactome visualised in **Figure S4D-G**). In comparison to only Mcl-1 redistribution or accumulation, substantially more tBid was required for Bax and Bak oligomerisation when both Mcl-1 accumulation and redistribution were taken into account (**Figure 4K**). Defining a threshold of 10% of Bax and Bak in pores as a minimum for efficient MOMP, both processes jointly elevated MOMP resistance approx. 2-3 fold from G_1_ to M phase (**Figure 4K,L**).

Taken together, these simulation results suggest that both accumulation and redistribution contribute substantially to apoptosis resistance and that both processes are required to achieve peak MOMP resistance late in the cell cycle.

### Mcl-1 subcellular localization and expression levels independently contribute to regulating apoptosis susceptibility

Next, we set out to study if Mcl-1 redistribution and Mcl-1 accumulation indeed independently contribute to elevated apoptosis resistance in experiments. We first measured the heterogeneity of Mcl-1 expression levels and subcellular distributions in H460-S-Mcl-1 cells. To make these cells dependent on endogenous Mcl-1 for survival, we inhibited both Bcl-2 and Bcl-xL (10 µM ABT-199, 10 µM WEHI-539), and then analysed the relationship between single-cell Mcl-1 abundance and localization, and the kinetics of cell death upon Mcl-1 inhibition (1 µM S63845) (**Figure 5A**).

**Figure 5:**
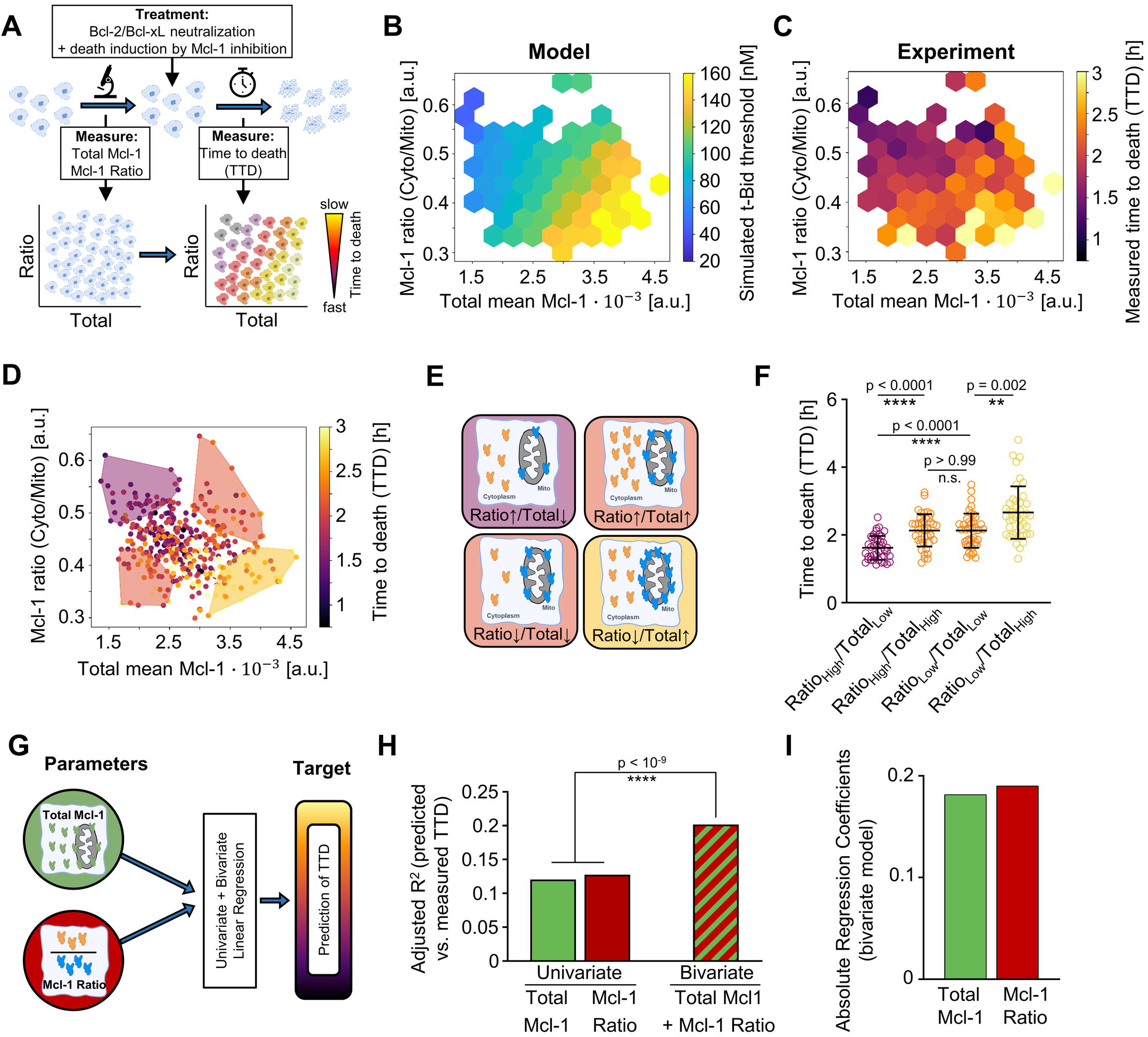
Mcl-1 subcellular localization and expression levels independently contribute to regulating apoptosis susceptibility. A) Experimental scheme. Living H460-S-Mcl-1 cells were measured on stage for their Mcl-1 expression and distribution. Cells were then treated with BH3 mimetics, with Mcl-1 inhibition driving cell death. Individual times to death were recorded. B) Hexbin plot showing the predicted t-Bid threshold of 420 cells pooled from 3 independent, batch-corrected experiments. t-Bid thresholds are color-coded and plotted against the respective Mcl-1 ratio and Mcl-1 total expression. C) Hexbinplot showing the time to death (TTD) of 420 cells pooled from 3 independent, batch-corrected experiments. TTD is color-coded and plotted against the respective Mcl-1 ratio and Mcl-1 total expression. D) 10% of cells in each corner were grouped and the average time to death of each group is depicted as a colored area. E) Scheme depicting the Mcl-1 expression and distribution in the four groups. F) Individual cells of the four groups were compared using One Way ANOVA with Dunnett’s T3 multiple comparisons. G) The contribution of either the ratio or the total Mcl-1 to the times to death was determined by a comparative approach using univariate linear regressions and multivariate linear regression via ANOVA. H) The adjusted R^2^ values of predicted vs. measured times to death from either the univariate regressions or the bivariate regression are depicted and compared using ANOVA and F-statistic. I) Absolute regression coefficients of the bivariate model are depicted.

To sufficiently capture population heterogeneity, we first measured Mcl-1 expression and distribution in a total of 420 cells and then applied our mathematical model to estimate MOMP resistances of these cells (**Figure 5B**). As would be expected from independent contributors to cell death resistance, MOMP thresholds increased with Mcl-1 expression and also with its redistribution towards the mitochondria (**Figure 5B**). Experimentally measured apoptosis sensitivities (time to death (TTD) after Mcl-1 inhibition) indeed corresponded very well to this prediction (**Figure 5C**), and both Mcl-1 expression and mitochondrial accumulation correlated with survival times (**Figure S5A,B**). For a more thorough and quantitative analysis of the effects of Mcl-1 expression and distribution on survival times, we compared cells with similar Mcl-1 expression but different distributions, or vice versa (**Figure 5D,E**). While cells with a high Mcl-1_cyto/mito_ ratio and low Mcl-1 expression died fastest, either a redistribution towards mitochondria or an increase in overall Mcl-1 expression was sufficient to significantly extend survival times (**Figure 5F**). Cells with high Mcl-1 expression and high accumulation at the mitochondria survived the longest (**Figure 5F, Figure S5C**). We further examined whether other factors, such as cell size or cell positioning, additionally influenced the measured survival times. However, these parameters showed only minimal explanatory power, compared to the distribution of Mcl-1 and overall Mcl-1 expression (**Figure S5D**).

To quantify the relative contributions of Mcl-1 distribution and expression on the survival times, we performed univariate and bivariate linear regressions (**Figure 5G**). We found, that the bivariate model was significantly better in predicting survival durations (TTD) than the individual univariate models (**Figure 5H**), which confirms that both parameters contribute significantly to apoptosis sensitivity. Importantly, the absolute values of the regression coefficients for overall Mcl-1 amounts and its distribution ratios were within a similar range, demonstrating from experimental data that both parameters contribute equally strong to apoptosis resistance (**Figure 5I**).

Taken together, these experimental findings demonstrate that the Mcl-1 distribution ratio and total Mcl-1 expression levels act independently of one another and contribute to apoptosis resistance to a comparable degree.

### Patient tumor heterogeneity in Mcl-1 distributions contributes to high cell-to-cell variability in expected MOMP resistance

Our results show that proliferating cancer cells cycle through phases of differential apoptosis resistance, driven by changes in both the abundance and subcellular localisation of Mcl-1. This dynamic regulation creates substantial cell-to-cell heterogeneity in susceptibility to MOMP. We next asked whether similar heterogeneity could also be observed in human tumour tissues.

To address this, we evaluated treatment-naïve stage III colorectal cancer tissue samples that had previously been analysed by multiplex immunofluorescence imaging of apoptosis signalling proteins (Lindner et al., 2022). In brief, stained samples from four patients underwent cell segmentation to achieve single-cell resolution, followed by cell type assignment into stromal, immune and tumour compartments. In the present study, we extended this analysis to include subcellular segmentation into cytoplasmic and mitochondrial regions, quantitative assessment of Mcl-1 expression and distribution, and model-based calculation of MOMP resistance for individual cells (**Fig.6A**). Individual cells were segmented based on nuclear DNA and cytoplasmic S6-kinase staining. Mitochondria were segmented using a combination of Bak and Smac signals (**Figure 6B**). Tumour cells were identified using the epithelial markers AE1 and PCK26 (**Figure 6C**, **Figure S6A**). As expected, mitochondrial compartments exhibited elevated amounts of Mcl-1 when compared with nuclear and cytoplasmic regions (**Figure S6B**). Moreover, Mcl-1 distribution ratios in tumour cells closely matched those observed in human cancer cells (**Figure S6C**). Both total Mcl-1 expression levels and its subcellular distribution displayed pronounced heterogeneity across all four patient samples (**Figure 6D**), comparable to the heterogeneity observed in our cell line experiments.

**Figure 6:**
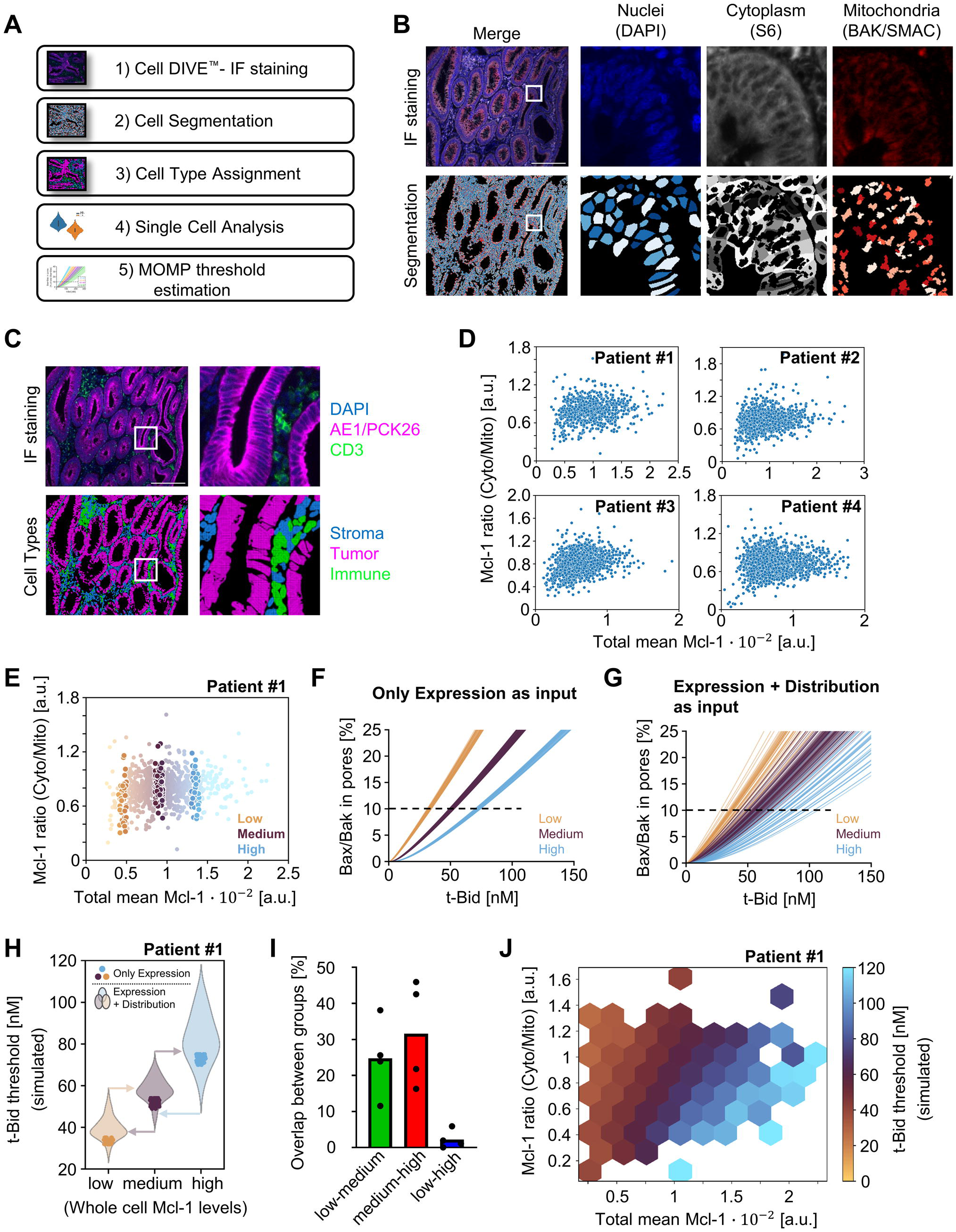
*In vivo* heterogeneity in Mcl-1 distributions contributes to high cell-to-cell variability in expected MOMP resistance. A) Scheme depicting the workflow of segmentation and analysis of the tumour samples. B) IF images of the compartment markers DAPI (for nuclei), S6-Kinase (for cytoplasm) and combined Bak/Smac signals (for mitochondria) with the respective masks. C) Classification of single cells into stroma, tumour or immune cells based on tumour markers AE1/PCK26 and immune marker CD3. Only tumour cells were used for further analysis. D) Mcl-1 distribution ratios and Mcl-1 expression of single tumour cells from four different samples. E) Cells from each tumour sample were classified into three groups based on their single-cell Mcl-1 expression (low, medium or high). F) The mean Mcl-1 expression of the cells grouped in (E) was used as input for the MOMP susceptibility model to simulate t-Bid sensitivity and Bax/Bak pore formation. G) The mean Mcl-1 expression as well as the subcellular Mcl-1 distribution of the cells grouped in (E) was used as input for the apoptosis susceptibility model to simulate t-Bid sensitivity and Bax/Bak pore formation. H) The t-Bid threshold values were compared between the groups. Thresholds with only Mcl-1 expression levels as input are depicted as solid filled dots. Thresholds derived from combined Mcl-1 expression and distribution data as inputs are depicted as violin plots. I) The percentage overlap between the groups was calculated for each patient sample. J) Hexbinplot showing the simulated t-Bid thresholds of single tumour cells dependent on the Mcl-1 distribution ratio and total Mcl-1 expression. Thresholds increase along both axes.

We next investigated the extent to which heterogeneity in Mcl-1 distribution impacts predicted MOMP resistance in tumour cells. First, we estimated MOMP resistance based solely on total Mcl-1 abundance, analogous to classical pathological stratification into low, medium, and high expressing samples, but at single-cell resolution (**Figure 6E**). When simulating the apoptosis sensitivity of these cells, increasing Mcl-1 abundance expectedly correlated with increased MOMP resistance (**Figure 6F**). Importantly, incorporating the measured subcellular distribution of Mcl-1 into the model markedly increased the heterogeneity in predicted MOMP resistance within each expression group (**Figure 6G**). Consequently, cell populations that appeared well separated based on Mcl-1 abundance alone now substantially overlapped in their predicted MOMP resistance (**Figure 6H, I; Figure S6D**). Extending this analysis beyond the discrete low, medium, and high expression groups to all analysed cells produced a very similar pattern to that observed in cell lines: both Mcl-1 abundance and its distribution between subcellular compartments contribute to determining the overall MOMP threshold (**Figure 6J, Figure S6E**). In other words, cells with similar total Mcl-1 levels are expected to differ markedly in their apoptosis sensitivity due to differences in the subcellular distribution of Mcl-1.

Taken together, these findings demonstrate that both Mcl-1 expression levels and intracellular localisation influence the predicted apoptosis sensitivity of individual tumour cells, with both the amounts and distributions independently contributing to high heterogeneity in expected apoptosis resistance within human tumour tissue.

## Discussion

In this study, we show that Mcl-1 expression and its redistribution between cytoplasm and mitochondria are coordinated with cell cycle progression. Both processes function as parallel, independent and equally potent rheostats that increase apoptosis resistance as cells progress through the cell cycle.

While cell cycle dependent regulation of Mcl-1 abundance has been reported before at population and single-cell scales (Harley et al., 2010; Pollak et al., 2021), we here, for the first time, identify Mcl-1 redistribution as an independent process and quantitatively define its impact on apoptosis resistance. Our results indicate that Mcl-1 exists in both soluble pools and mitochondria-associated pools, the latter including tightly bound or membrane-integrated fractions. The existence of different Mcl-1 pools is supported by reports identifying Mcl-1 fractions in the nucleus and cytosol (Fu et al., 2022; Kale et al., 2018; Nakajima et al., 2014) and by evidence for fractions of Mcl-1 engaging in different interactions at and within mitochondria. For example, pools of Mcl-1 were reported to localise to the mitochondrial matrix, where they affect mitochondrial respiration and fusion (Perciavalle et al., 2012). Mcl-1 can also associate with ACSL1 in the outer mitochondrial membrane to regulate long-chain fatty acid oxidation (Wright et al., 2024), and Mcl-1 obviously can engage other Bcl-2 family members in the outer mitochondrial membrane (Kale et al., 2018). Given the rapid exchange of the mobile Mcl-1 pool between mitochondria and cytosol identified here, it is tempting to speculate that a substantial pool of Mcl-1 is membrane associated rather than integrated, thereby permitting swift exchange. Dynamic shuttling of other Bcl-2 family members, most notably during Bcl-xL dependent Bax and Bak retrotranslocation from mitochondria into the cytosol, has been reported before (Edlich et al., 2011; Todt et al., 2015). However, the cell cycle dependent regulation of Mcl-1 localization appears distinct, as we did not observe a comparable regulation for Bcl-xL. Moreover, Mcl-1 exchange kinetics are remarkably fast, exceeding both the reported rates of Bcl-xL mediated retrotranslocation and Mcl-1 protein turnover (Edlich et al., 2011; Nijhawan et al., 2003; Slomp et al., 2021; Todt et al., 2015).

The accumulation of Mcl-1 on mitochondria from S phase and peaking in M phase depends on both increased Mcl-1 abundance and its net redistribution towards mitochondria. Cells also increase in volume and mitochondrial content as they proceed through the cell cycle (Posakony et al., 1977), which could contribute to higher local Mcl-1 levels. In both our simulations and measurements, we corrected for these factors, yet still observed a net accumulation of Mcl-1 on mitochondria. This indicates that redistribution must be driven by other processes that promote translocation to mitochondria, reduce retrotranslocation, or both.

We show that daughter cells initially inherit the peak Mcl-1 amounts achieved in their mother cell during M phase, before these Mcl-1 amounts are reset back to the lower levels typical of G_1_ phase cells. This may appear to contradict previous studies reporting that Mcl-1 degradation begins during M phase (Allan et al., 2018; Clarke et al., 2018; Haschka et al., 2015; Wertz et al., 2011). However, those studies largely focused on scenarios of prolonged mitotic arrest and made use of chemically synchronized cell populations. It can be conceived that under prolonged arrest, degradation mechanisms are activated that are absent, or activated later, during or after unperturbed mitosis. Moreover, chemical cell synchronization can substantially influence cell cycle- and proliferation-associated processes (Cooper, 2003; Ligasová & Koberna, 2021; Min et al., 2020), potentially interfering with the dynamics studied here. Still, we cannot formally exclude the possibility that enhanced Mcl-1 degradation is at least initiated in M phase, without notably affecting protein amounts yet. Since cytokinesis strongly reduces daughter cell volumes compared with the mother cell, Mcl-1 concentrations could possibly be maintained initially despite ongoing degradation. Importantly, also the distribution of Mcl-1 is inherited to daughter cells, so that newly divided cells are expected to initially exhibit high apoptosis resistance. Indeed, our prior work on transmitotic cell fates during extrinsic apoptosis showed that daughter cells inheriting high Mcl-1 levels and active caspase-8 from mother cells require at least 30 to 60 min to regain MOMP competency (Pollak et al., 2021).

While both Mcl-1 amounts and localisation are regulated across the cell cycle, we observed substantial cell-to-cell heterogeneity in both parameters, even among cells from the same cell cycle stage. Because we could show that localisation is as important for cell fate decision making as overall Mcl-1 amounts, this represents an independent and equally significant layer contributing to cell-to-cell variability in apoptosis sensitivity. Although non-genetic heterogeneity in cell death susceptibility is naturally revealed in single cell studies (Rehm et al., 2002; Spencer et al., 2009), it is now increasingly recognized as a critical contributor that can allow some cells to evade death and potentially drive cancer relapse and disease progression (Ichim et al., 2015; Russo et al., 2024). Our analyses of patient tumour samples confirm that heterogeneity in Mcl-1 expression and subcellular distribution also occurs among cells from individual tumours. Moreover, our estimates of apoptosis thresholds indicate that both parameters strongly influence expected cell death resistance, even blurring distinctions between cell populations that would be clearly separable based on Mcl-1 expression alone.

In our study, we focussed on identifying the dynamics of spatiotemporal Mcl-1 regulation and their relevance for functional consequences affecting cell fates. Our evidence that Mcl-1 spatial regulation significantly affects cellular life/death decisions, experimentally shown to be as important as overall Mcl-1 accumulation, provides a justification for future studies aimed at dissecting the likely complex and multifactorial mechanisms controlling Mcl-1 subcellular localization. These mechanisms might be linked to Mcl-1 itself, its interaction partners, or features of mitochondria that change over the cell cycle and specifically affect Mcl-1 rather than Bcl-xL. Because Mcl-1 pools with distinct mobilities exist, understanding these underlying processes could also enable the development of pharmacological strategies that selectively affect the subcellular localization of specific Mcl-1 pools. This might contribute to novel therapeutic avenues by which systemic toxicities encountered upon global Mcl-1 inhibition could be avoided (Bolomsky et al., 2020; Kelly & Strasser, 2020).

## Materials and methods

### Reagents and antibodies

Primary antibodies were as follows: mouse anti-α-Tubulin (DM1A, Cell Signaling #3873, 1:1000 western blotting (WB)), rabbit anti-Calnexin (Cell Signaling #2433, 1:1000 WB), rabbit anti-Cox IV (3E11, Cell Signaling #4850, 1:1000 WB), mouse anti-GAPDH (D4C6R, Cell Signaling # 97166, 1:1000 WB), mouse anti-Lamin A/C (Cell Signaling #4777, 1:1000 WB), rabbit anti-Mcl-1 (D2W9E, Cell Signaling # 94296, 1:1000 WB, 1:800 IF, 1:50 flow cytometry (FC)), mouse anti-RFP (ChromoTek 6g6, 1:1000 WB), rabbit anti-Phospho-Histone H3 (D2C2, Cell Signaling #3377 and #8552, 1:50 FC), rabbit anti-Tom20 (Cell Signaling #42406, 1:1000 WB).

Secondary antibodies were as follows: goat anti-rabbit IgG – Alexa Fluor 647 (Life Technologies Corporation), goat anti-rabbit IgG – Alexa Fluor 488 (Life Technologies Corporation), goat anti-rabbit IgG – Peroxidase (Dianova GmbH), goat anti-mouse IgG – Peroxidase (Dianova GmbH), goat anti-rabbit IgG - IRDye®680RD (LI-COR GmbH), goat anti-mouse IgG - IRDye®800CW (LI-COR GmbH).

Fc-scTRAIL was produced as described previously (Hutt et al., 2017). DMSO was purchased from Carl Roth, ABT-199 from Active Biochem, S63845 and WEHI-539 from APExBIO and bortezomib from UBPBio. MioTracker™ Red CMXRos, MitoTracker™ Green FM and DAPI were obtained from Invitrogen, cycloheximide was purchased from Sigma. Puromycin was obtained from AppliChem GmbH.

### Plasmids and transfections

Plasmid EZ-Tet-pLKO-Blast (Addgene #85973) was used to express siRNA against Mcl-1 (CCAGTATACTTCTTAGAAAGT), cloned as a hairpin sequence for stabilization (CTAGC-CCAGTATACTTCTTAGAAAGT-TACTAGT-ACTTTCTAAGAAGTATACTGG-TTTTT-G). This plasmid was lentivirally transduced in NCI-H460 cells.

Plasmids pSpCas9(BB)-2A-Puro human MCL-1 and pUC Scarlet-MCL-1 were both received from Stephen Tait (Cao et al., 2022) and used to endogenously tag Mcl-1 with mScarlet in NCI-H460 cells. Silencer®Select siRNA targeting Mcl-1 (ambion #s8583; CCAGUAUACUUCUUAGAAATT) was transfected using Lipofectamine®RNAiMAX (Thermo Fisher Scientific) according to the manufacturer’s protocol.

### Tagging endogenous Mcl-1 with mScarlet

NCI-H460 cells were seeded in a 10 cm petri dish and transfected at roughly 30% confluence. pSP-Cas9 and pUC-mScarlet plasmid DNA (kindly received from Stephen Tait, published in (Cao et al., 2022), was prepared with the Lipofectamine 3000 transfection kit (Thermo Fisher) according to the manufacturer’s protocol and added onto the cells. 24 hours after transfection, the medium was changed to medium containing 0.5 µg/ml puromycin for the selection of transfection-positive cells. Cells were sorted at the FACS for their mScarlet fluorescence and expanded as single cell clones. The mScarlet intensity of the expanded clones was measured via flow cytometry with excitation at 561 nm and a 586/15 nm emission filter. The molecular size of the mScarlet-Mcl-1 construct was evaluated via immunoblot for Mcl-1 and mScarlet. The subcellular distribution of mScarlet-Mcl-1 was validated using live cell imaging at the LSM 980 microscope. To validate the mScarlet insertion, genomic DNA was amplified using the quick extract DNA kit (Biozym, #QE0905T) according to the manufacturer’s protocol. The mScarlet-insertion was amplified via PCR and the PCR products were successfully sequenced and confirmed a correct knock-in. The following primers were used to amplify the Mcl-1 locus: Mcl-1-UTR-for: CACTTCCGCTTCCTTCCAGT; Mcl-1-Ex1-rev2: CCGCGTTTCTTTTGAGGCCA.

PCR was done using the PCR-NEB Q5 Kit. PCR samples were loaded on an 1% agarose gel and DNA was stained via EtBr in the gel. Respective bands were cut out of the gel and sequenced.

### Cell culture

Cells were cultivated with RPMI 1640 medium, supplemented with 10% (v/v) FBS in cell culture flasks (37°C, 5% CO_2_). Cells were seeded and passaged with a passaging ratio between 1:3 and 1:10.

### Immunofluorescence imaging and image analysis

To analyse subcellular protein concentrations, cells grown on coverslips were incubated with MitoTracker Red CMXRos (100 nM) for 90 minutes prior to fixation with 4% PFA, permeabilization with 0.3% Triton-X-100 and immunostaining. The nuclei were stained using DAPI. Images were acquired on a Zeiss Axio Observer SD Spinning Disk microscope equipped with a PlanApochromat 40×/1.4 NA oil objective and an Axiocam 503 Mono CCD camera. Geminin was excited with a 488 nm diode laser using a 525/50 nm emission filter, the Alexa Fluor 647 dye was excited with a 638 nm diode laser using the 690/50 nm emission filter and DAPI was excited with a 405 nm diode laser using the 450/50 nm emission filter. MitoTracker Red was excited with a 561 nm diode laser using a 575/50 nm emission filter. All images were taken as Z-stacks with 0.5 µm distance between each Z-layer. For quantification, a single Z-layer close to the surface of the coverslip was analysed using CellProfiler 4.2.7 (Stirling et al., 2021). In the CellProfiler analysis, nuclei were recognized through DAPI staining and the respective cell bodies were expanded around each nucleus based on the Mcl-1 staining. Based on the MitoTracker signal, the mitochondrial area was defined for each cell individually. The cytoplasm represents the inverse of the mitochondria, the nucleus was defined as a separate object. To quantify subcellular protein expression, the respective IF staining intensities were measured. The distinction between early and late cell cycle stages was set stringently at the 0.15 and 0.85 percentile of geminin mean intensities inside the nucleus, cells in between were not considered for cell cycle specific analysis. For the specific analysis of M cells and newly divided sibling cells, DNA- and cell morphology were manually evaluated and categorized.

### Time-lapse imaging and image analysis

For analysis of transmitotic apoptosis resistance, cells were plated on 35 mm glass-bottom dishes (CellView Cell Culture Dish, Greiner Bio One) in Phenol Red-free RPMI 1640 containing 10% FBS. Images were acquired at 37°C and 5% CO_2_ on a Zeiss Cell Observer microscope equipped with an Axiocam MRm CCD camera and a Plan-Apochromat 20×/0.8 objective. Medium containing Fc-scTRAIL alone or in combination with S6385 was added and cells were imaged for 24 h in 15 minute intervals. The time until death (t_death_) after treatment was measured for individual cells. Thereby, the population of analysed cells was subdivided into cells that underwent mitosis (f0+f1) or not (f0).

For analysis of Mcl-1 expression and distribution, cells containing endogenous-tagged mScarlet-Mcl-1 were plated on 35 mm glass-bottom dishes (CellView Cell Culture Dish, Greiner Bio One) in Phenol Red-free RPMI 1640 containing 10% FBS. Before imaging, cells were incubated with Phenol Red-free RPMI 1640 containing 10% FBS and 100 nM MitoTracker Green and 1 µg/ml Hoechst 33342 for 30 minutes at 37°C and 5% CO_2_. Images were acquired at 37°C and 5% CO_2_ on a confocal laser scanning microscope (LSM 980 Airyscan 2) equipped with a Plan-Apochromat 63×/1.40 Oil DIC M27 objective. mScarlet-Mcl-1 was excited with a 561 nm laser using a 573-627 nm emission filter, MitoTracker Green was excited with a 488 nm laser using a 490-512 emission filter. If added to the experiment, Hoechst was excited with a 405 nm laser using a 380-548 emission filter. Cell segmentation was done as described for immunofluorescence images, using the Hoechst staining for nuclei, mScarlet-Mcl-1 signal for the cell body and MitoTracker Green for mitochondria. For the Fluorescence Loss in Photobleaching (FLIP) experiments, measurement regions and control cells were manually cropped and analysed via CellProfiler as describe above.

### Flow cytometry

Cells were harvested and centrifuged at 300 *g* and 4°C for 5 minutes. The supernatant was aspirated and the pellet was resuspended in 100 µl 4% PFA in PBS for 20 minutes at RT. After PFA fixation, the cells were centrifuged again at the same conditions as before and resuspended in 100 µl permeabilization solution 2 (BD Biosciences, Germany) for 20 minutes at RT. Followed by another centrifugation step, the cells were resuspended in 100 µl medium for at least 30 minutes at RT, to block unspecific binding sites. After centrifugation, desired target antigens were then primed by incubation with the respective primary antibody in medium for 90 minutes at RT, including an isotype control antibody. After another centrifugation step, cells were washed twice in PBS for 5 minutes each, including centrifugation in between. To detect the bound primary antibodies, the cells were incubated with fluorescently-tagged secondary antibodies. Incubation with the secondary antibody was done in RPMI 1640 medium for 45 minutes at RT. Following a final centrifugation step, the cells were resuspended in PBS supplemented with 0.02% (w/v) sodium azide and transferred into a 96-well plate for flow cytometry (MACSQuant VYB, Miltenyi Biotec, Germany).

### Western blotting

Cells were lysed in solubilization buffer [50 mM Tris-HCl pH 7.5, 150 mM NaCl, 1 mM EDTA, 1% (v/v) TritonX-100 plus Complete Protease Inhibitors (Roche) and 1 mM DTT], incubated on ice for 20 min and centrifuged at 21,000 *g* and 4°C. Protein concentrations were determined by Bradford assay (Roti®-Quant). Afterwards, samples were incubated in Laemmli buffer for 10 minutes at 98°C. Proteins were separated using Bolt^TM^ 4 – 12% Bis-Tris precast gels (Thermo Fisher) and blotted on a nitrocellulose membrane using the iBlot2® (Thermo Fisher). Unspecific binding sites on the membrane were blocked by incubation with western-blocking-reagent (Roche) 1:10 in TBS-T for at least 60 minutes at RT. Incubation with primary antibodies in PBS supplemented with 0.02% (w/v) sodium azide was done at 4°C overnight. Followed by three washing steps with TBS-T, the membrane was incubated with the a HRP-conjugated secondary antibody for 45 minutes at RT in TBS-T and 0.5X blocking reagent. After three more washing steps with TBS-T, the membrane was incubated with immobilon forte substrate (manufacturer) for 5 minutes at RT in the dark, followed by measurement at an Amersham™ imager 600 (GE Healthcare, USA). Alternatively, a fluorescently-labelled secondary antibody was used and imaged at a Li-Cor 9120 Odyssey® imager (LI-COR Biosciences GmbH, Germany).

### Subcellular fractionation

NCI-H460 were harvested and homogenized in ice cold homogenization buffer [225 mM mannitol, 75 mM sucrose, 0.1 mM EGTA/EDTA, 30 mM Tris-HCl pH 7.4] supplemented with cOmplete protease inhibitors. All subsequent centrifugation steps were performed at 4°C. To isolate the nuclear fraction, the homogenate suspension was centrifuged at 600 *g* for 5 min. The resulting pellet, containing nuclei and cell debris, was resuspended in homogenization buffer with 100 µg/ml digitonin for further purification and centrifuged again at 600 *g* for 5 min. The pellet was then resuspended in PBS, washed, and centrifuged once more at 600 *g* for 5 min to obtain the final nuclear pellet. The purified nuclei were stored at -20°C for subsequent analyses. The supernatant from the first centrifugation was centrifuged again at 600 *g* for 5 min to remove residual nuclei. The resulting supernatant was then centrifuged at 7,000 *g* for 15 min to obtain the mitochondrial fraction. The mitochondrial pellet was resuspended in 2 ml of ice-cold PBS, centrifuged again at 7,000 *g* and subsequently at 10,000 *g* for 10 min. The final mitochondrial pellet was stored at -20°C for further analysis. The supernatant from the 7,000 *g* centrifugation was further centrifuged at 20,000 *g* for 30 min. The remaining supernatant was then centrifuged at 100,000 *g* for 1 h. The resulting pellet contained the endoplasmic reticulum (ER), while the supernatant represented the cytosolic fraction. The cytosolic fraction was additionally precipitated by adding trichloroacetic acid (TCA) to a final concentration of 15%, followed by incubation on ice for 30 min. After centrifugation at 18,000 *g* for 10 min, the resulting pellet was washed by gently adding 100% ethanol. This step was repeated once. Proteins from the cytosolic fraction were dissolved in Laemmli buffer, heated at 95°C for 10 min, and analysed by Western Blot. For all other fractions, protein concentrations were first determined using the Bradford assay before further processing.

### Sodium carbonate extraction

Cells were harvested, centrifuged at 400 *g* for 5 min. and the pellet was resuspended in 5 ml homogenization buffer containing cOmplete protease inhibitors and kept on ice for 2 min. The cells were then mechanically homogenized with a glass pestle by gently moving it up and down. Successful homogenization was confirmed under a microscope. To remove intact cells, the suspension was centrifuged at 3,200 *g* and 4°C for 5 min. The supernatant was collected, and the centrifugation step was repeated until no sediment was observed. The supernatant was divided into four aliquots and centrifuged at 18,000 *g* and 4°C for 10 min. The resulting pellet was resuspended in 1 ml homogenization buffer and centrifuged again at 18,000 *g* and 4°C for 10 min. The pellets containing the crude mitochondria fraction were resuspended in 100 mM Na_2_CO_3_ with pH values of 10, 11.25 and 12.5, or in PBS as a no treatment control, and incubated on ice for 30 min. The samples were centrifuged at 100,000 *g* and 4°C for 60 min. The insoluble fraction in the pellet was dissolved in Laemmli buffer and heated at 98°C for 10 min. The soluble fraction in the supernatant was precipitated by adding TCA. The final samples were analysed using immunoblotting.

### Analysis and quantification of Mcl-1 shuttling kinetics via FLIP

In each cell, a bleaching area at one edge of the cell, and a measurement area at the opposing edge of the same cell was defined. After two minutes of equilibration, mScarlet-Mcl-1 was bleached within the bleaching area using 100% power of the 561 nm laser every 10 seconds. Simultaneously, the decay of mScarlet intensity in the measurement area was measured every 10 seconds. The measurement area was segmented into mitochondrial and cytoplasmic compartments based on MitoTracker green staining. To correct for unwanted bleaching through light dispersion, a control cell next to the FLIP cell was measured in the same way. A one-phase exponential decay function 𝑓(𝑡) was fitted to the mScarlet-Mcl-1 decay intensities 𝐼(𝑡) of the control cell, to quantify unspecific bleaching outside of the bleaching area. Thus, the raw data 𝐼(𝑡) was corrected by dividing it by 𝑓(𝑡) and scaling to 𝑓(0), which led to the corrected control decay 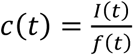· 𝑓(0). In the same way, the data from the FLIP cell was corrected using the corrective function 𝑓(𝑡) from the control cell on the FLIP data. To obtain kinetic parameters of Mcl-1 (𝑘_𝑜𝑛_ or 𝑘_𝑜𝑓𝑓_), a simplified ODE model was defined that described the FLIP experiment. The model consisted of a cytoplasmic and a mitochondrial compartment and a theoretical external compartment. Mcl-1 can be either localized at the mitochondria or in the cytoplasm, or existing in the external compartment as “bleached Mcl-1”. In the model, the bleaching was simulated by depletion of cytoplasmic Mcl-1 into the external compartment. Mcl-1 was assumed to bind to the mitochondria and be released back into the cytoplasm based on mass-action kinetics. This model was mathematically written as a system of three coupled ODEs:

Equation (I) – ODE of cytoplasmic Mcl-1 in the FLIP model:

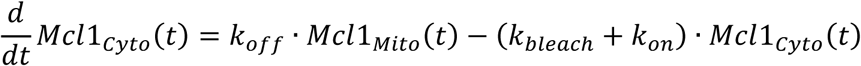

Equation (II) – ODE of mitochondrial Mcl-1 in the FLIP model:

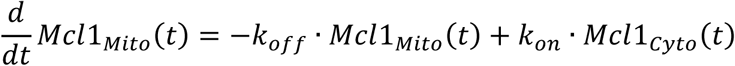

Equation (III) – ODE of bleached Mcl-1 in the FLIP model:

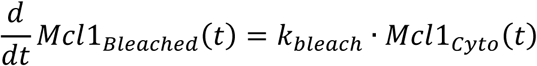

Using the SymPy library in python, this ODE system was solved analytically to efficiently optimize the kinetic parameters of the system to the experimental data. Therefore, the initial conditions 𝑀𝑐𝑙1_𝐶𝑦𝑡𝑜_(0) and 𝑀𝑐𝑙1_𝑀𝑖𝑡𝑜_(0) were taken from the experimental data and 𝑀𝑐𝑙1_𝐵𝑙𝑒𝑎𝑐ℎ𝑒𝑑_(0) was set to zero. For each optimization step, the ODE system was evaluated every 10 s for 10 minutes, similar to the experiment. Then, an exponential decay function was fitted through these modelled decay curves of 𝑀𝑐𝑙1_𝐶𝑦𝑡𝑜_ and 𝑀𝑐𝑙1_𝑀𝑖𝑡𝑜_ and compared to the measured decay curves in the respective compartments. The loss was defined as the absolute difference in the decay coefficients between the ODE model and experimental data. Using the DIviding RECTangles (DIRECT) optimization algorithm from the SciPy library, a pre-defined parameter space of the kinetic parameters was iteratively screened for a global parameter optimum that fitted the experimental data.

### Plasma membrane permeabilization using digitonin

Cells were stained with MitoTracker for 90 minutes. Prior to fixation, cells were treated with 100 µg/ml Digitonin in PBS for 1 min at RT, and washed of immediately after with PBS. After permeabilization, the cells were kept in PBS for different time points before fixation and immunofluorescence staining.

### In silico cell cycle model of Mcl-1 expression and distribution

The ODE model consisted of a volumetric mitochondrial and cytoplasmic compartment, that contain mitochondrial or cytoplasmic Mcl-1, respectively. Therefore, it was implemented within the python library Tellurium in Antimony (Choi et al., 2018; Smith et al., 2009).

Mcl-1 was assumed to shuttle between the cytoplasmic and the mitochondrial compartments obeying the experimentally defined distribution ratio and the shuttling rates defined by FLIP measurements:

Equation (IV) – ODE of cytoplasmic Mcl-1 in the cell cycle model:

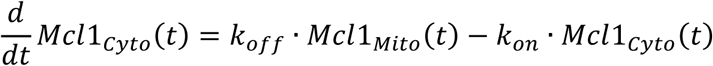

Equation (V) – ODE of mitochondrial Mcl-1 in the cell cycle model:

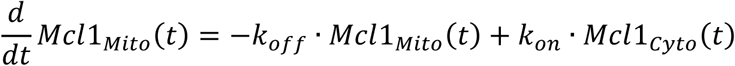

To include the cell cycle, the lengths of the cell cycle phases (G_1_, combined S/G_2_ and M) were determined using flow cytometry and respective cell cycle phase markers. By dividing the total cell cycle length of 18 h by the respective percentages of cells in each phase, the duration of each phase was estimated. By using growth rates from literature (Cadart et al., 2022), the volume of the compartments was dynamically defined to exponentially double over the time of one cell cycle and was reset to starting values upon division. Since 𝑀𝑐𝑙1_𝐶𝑦𝑡𝑜_ and 𝑀𝑐𝑙1_𝑀𝑖𝑡𝑜_ were defined as concentrations, volume growth also influenced these values in the ODE system:

Equation (VI) – Dynamic change in mitochondrial volume:

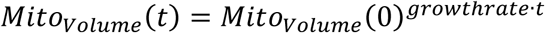

Equation (VII) – Dynamic change in cytoplasmic volume:

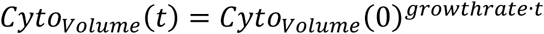

Mcl-1 turnover was implemented in the model by a continuous production of cytoplasmic Mcl-1 that exponentially increases with cell growth, since Mcl-1 mRNA concentrations was found to remain constant with cell cycle progression (Pollak et al., 2021). Mcl-1 degradation was defined based on mass-action law, with the degradation rate calculated from a Mcl-1 half-life of 30 minutes (Nijhawan et al., 2003; Slomp et al., 2021).

Equation (VIII) – Turnover addition to Equation (IV):

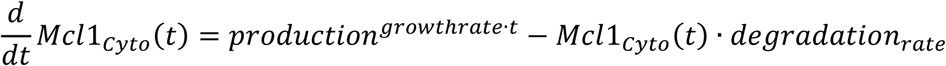

To account for the cell cycle dependent accumulation of Mcl-1, the degradation rate was exponentially decreased with cell cycle progression, with the rate empirically fitted to experimental data:

Equation (IX) – Dynamic decrease in degradation rate:

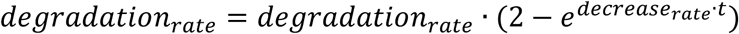

To account for the cell cycle dependent change in Mcl-1 distributions, the ratio between 𝑘_𝑜𝑛_ and 𝑘_𝑜𝑓𝑓_ was adjusted over time. Since 𝑘_𝑜𝑛_ and 𝑘_𝑜𝑓𝑓_ are directly dependent on each other through the ratio between cytoplasmic and mitochondrial Mcl-1, 𝑘_𝑜𝑛_ can be described as 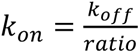. In the model, the ratio value was dynamically changed to match the experimental observations in early and late cell cycle stages.

The overall model was solved using the roadrunner library in python (Somogyi et al., 2015). This allowed to simulate any single cell or cell ensembles over any number of cell cycles based on the following initial parameters: Mcl-1 expression level, Mcl-1 distribution, Cell Size, Cell Cycle Stage.

For predictive perturbation simulations, Mcl-1 accumulation or redistribution were independently deactivated. Mcl-1 accumulation was deactivated by keeping the degradation rate constant over the whole cell cycle, which leaves Mcl-1 turnover in a steady equilibrium and constant concentration. Mcl-1 redistribution was deactivated by keeping the ratio value constant over the whole cell cycle.

### Time to death experiments and analysis

H460-S-Mcl1 cells were seeded and stained with 100 nM MitoTracker Green and imaged as mentioned above. Multiple fields of view with at least 10 cells each were defined and imaged to capture Mcl-1 expression and distribution. After the initial image, the cells were treated on stage with a BH3 mimetic combination of 10 µM ABT-199, 10 µM WEHI-539 and 1 µM S63845 in medium. Following the treatment, the cells were continuously imaged every 5 minutes to determine the time until each cell morphologically died (membrane blebbing). Single cell parameters such as mScarlet-Mcl-1 intensities, compartment sizes or cell positions were analysed from the initial image. For each cell, the Mcl-1 expression and Mcl-1 distribution were used as input for the above-mentioned apoptosis susceptibility model to simulate an individual t-Bid threshold. Further, correlations of Mcl-1 expression and distribution were analysed using Pearson’s correlation coefficient. To determine the extreme 10% per corner, the distances of each cell to each corner were calculated and the 10% of cells that were closest to each corner were assigned to that respective group. Univariate and bivariate regressions and the resulting R^2^ values and regression coefficients were analysed in python using the Scikit-learn library. Variance inflation factor analysis was done using the Statsmodels library in python.

### Colorectal cancer tissue samples and multiplex imaging

Formalin-fixed, paraffin-embedded primary tumor tissue sections were obtained from four chemotherapy-naïve, resected stage III CRC patients collected from Queen’s University Belfast (UK). Three samples from the core of each tumour were assembled on tissue microarrays (TMAs). Multiplexed immunofluorescence iterative staining of the TMAs was performed as previously described (Lindner et al., 2022) using the Cell DIVE™ technology (Leica Microsystems; formerly GE Healthcare). This involved iterative staining and imaging of the same tissue section with multiple antibodies and is achieved by mild dye oxidation between successive staining and imaging rounds. In total, there were 13 staining rounds using cell segmentation, cell identity and apoptosis signalling antibodies as described in (Lindner et al., 2022), and DAPI was imaged in each round. The Leica Bond (Leica Biosystems) was used for antibody staining. Commercially acquired antibodies underwent multi-step process of validation and dye conjugation as previously described (Lindner et al., 2022). Exposure times were set to fixed values for all images of a given marker.

### Single cell segmentation and analysis in tumour samples

Raw images of stained tumour tissues were obtained as described (Lindner et al., 2022). In the present study, nuclei and surrounding cytoplasmic areas of single cells from each TMA were segmented using the Cellpose library in python. This library contains pretrained models that recognize nuclei and whole cell. The *cyto2* pretrained model was used together with the DAPI staining for nuclei and S6 kinase staining for the cytoplasm. The segmented masks were imported into CellProfiler, where a combined image of Bak and Smac staining was used to segment mitochondria within the cytoplasmic mask. Thereby, the rescaled Bak signal was weighted 2/3 and the rescaled Smac signal was weighted 1/3 of the resulting image. Mitochondrial structures were segmented using adaptive thresholding, so that tiny structures were size-excluded to minimize false-positive mitochondrial masking.

Classification of cells into tumour, immune or stroma cells was done using CD3 (immune marker) and AE1/PCK26 (tumour markers) intensity distributions. Cells negative for all markers were assigned as stroma. All analysis of Mcl-1 amounts and distributions were conducted in the tumour cells.

### Statistical analysis

Statistical analysis was performed using PRISM 10 (GraphPad Software) and the following libraries in python: Statsmodels, Scikit-learn, SciPy, NumPy. Unpaired t-test with Welch’s correction was used in Figures 1D,F,G; S1E; 3D; S2D,E. One-Way ANOVA with Dunnett’s T3 multiple comparisons was used in Figures 2E,F; S3B, 4 J,L; 5F; S6B,C. Mann-Whitney test was used in Figures 2H,I. Linear regression was performed, and nested models were compared using ANOVA and the F-test in Figures 5H,I.

### Visualisations

Schematics in figures 3K and S2A were created with BioRender.com

## Acknowledgments

The authors acknowledge Prof Stephen Tait (Cancer Research UK Scotland Institute, Univ. of Glasgow) and Dr Joel Riley (Medical University of Innsbruck) for providing crucial plasmids and helpful discussions, and Cristina Jaus (University of Stuttgart) for technical assistance. The authors also acknowledge the Technology Platform “Cellular Analytics“ of the Stuttgart Research Center Systems Biology for their support & assistance, Ms Elizabeth McDonough (GEHC Technology and Innovation center) for running Cell DIVE analysis of CRC stage III samples, and Dr. Sanghee Cho for data processing.

## Data availability

Data are available from the authors.

## Code availability

All codes developed for this study are available at: https://doi.org/10.5281/zenodo.18185366

## Author Contributions

FK: data acquisition, analysis and interpretation, draft writing; NP, AB: data acquisition, analysis and interpretation, draft revision; BK: data acquisition, draft revision; FG, DBL: draft revision, funding acquisition; JP: draft revision, supervision, funding acquisition; MR: data interpretation, draft writing, supervision, funding acquisition.

## Ethics declarations

Raw data related to Fig.6 and Fig.S6 were used in line with FAIR data principles from a previously published study (Lindner et al., 2022). Tissues were supplied by the Queen’s University Belfast Department of Pathology with written consent provided by all patients and institutional ethical approval granted. Ethical approval for processing of samples and clinical data was also obtained by the Beaumont Hospital Research and Ethics Committee.

