## Supplemental Figures for "Spatiotemporal dynamics of Mcl-1 abundance and its influence on apoptosis susceptibility"

### Figure S1

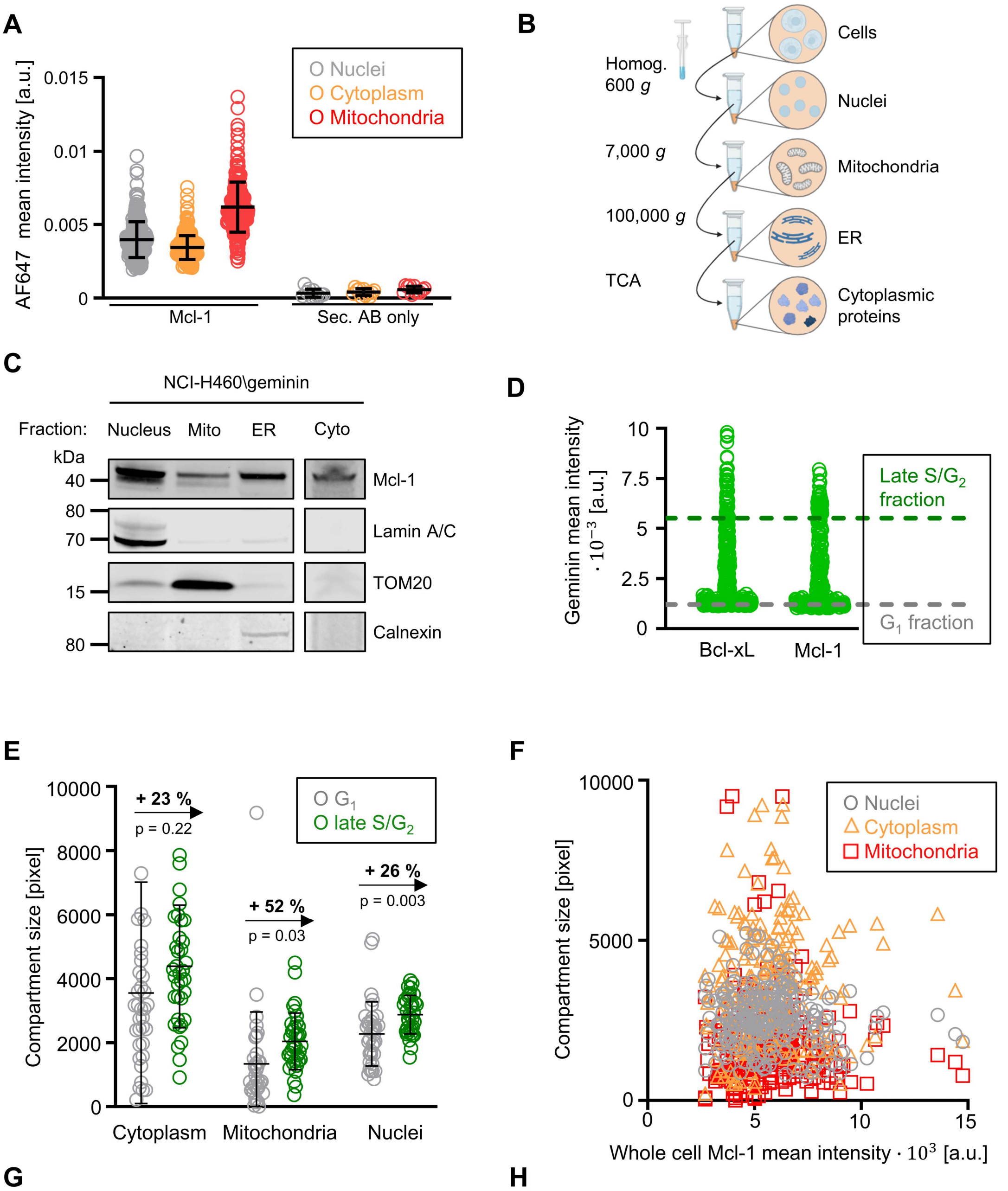


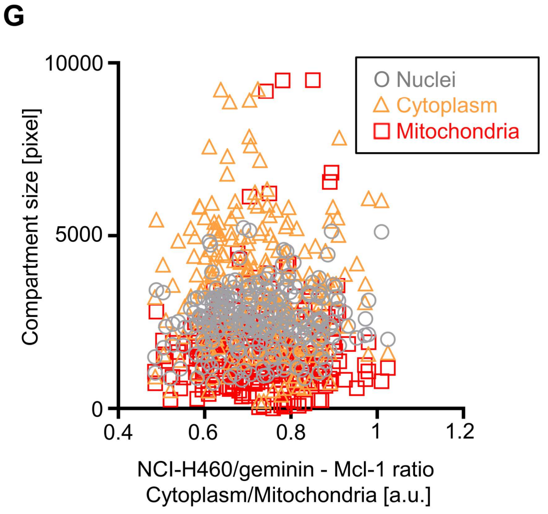


**H**

| Correlation n = 243 | Spearman r | *p* value |
|---|---|---|
| Nuclei size - Whole cell Mcl-1 | -0.08 | 0.2 |
| Cytoplasm size - Whole cell Mcl-1 | 0.07 | 0.26 |
| Mitochondria size - Whole cell Mcl-1 | 0.09 | 0.18 |
| Nuclei size - Mcl-1 ratio | 0.05 | 0.43 |
| Cytoplasm size - Mcl-1 ratio | -0.06 | 0.36 |
| Mitochondria size - Mcl-1 ratio | -0.03 | 0.69 |

**Figure S1**

A) Subcellular mean intensities of Mcl-1 were compared with background signals obtained by staining with only the secondary antibody.

B) Schematic overview of the subcellular fractionation workflow. After homogenization of the cells, organelle-enriched pellets were obtained by sequential centrifugation steps, and cytoplasmic proteins were isolated from the final supernatant by TCA precipitation.

C) Western blot analysis of Mcl-1 in subcellular fractions from NCI-H460/geminin cells. Blots were probed for Mcl-1 and compartmental marker proteins.

D) Geminin mean intensities of cells that were stained for either Bcl-xL or Mcl-1. Cells were separated into a $G_1$ fraction below the 0.15 percentile of geminin mean intensity and a late S/$G_2$ fraction above the 0.85 percentile, respectively.

E) Compartment sizes from the subcellular segmentation were measured and compared between the $G_1$ fraction and a late S/$G_2$ fraction. n = 35 cells per group.

F) Single cell compartment sizes were plotted against the whole cell mean Mcl-1 intensities.

G) Single cell compartment sizes were plotted against the Mcl-1 distribution ratios.

H) The correlations depicted in (F,G) were quantified by calculating Spearman's r and the associated *p*-value.

One representative experiment from at least three independent biological replicates is shown. Each dot represents a single cell. Error bars represent mean ± standard deviation (A,E). p-values from an unpaired t-test with Welch's correction are depicted in (E).

### Figure S2

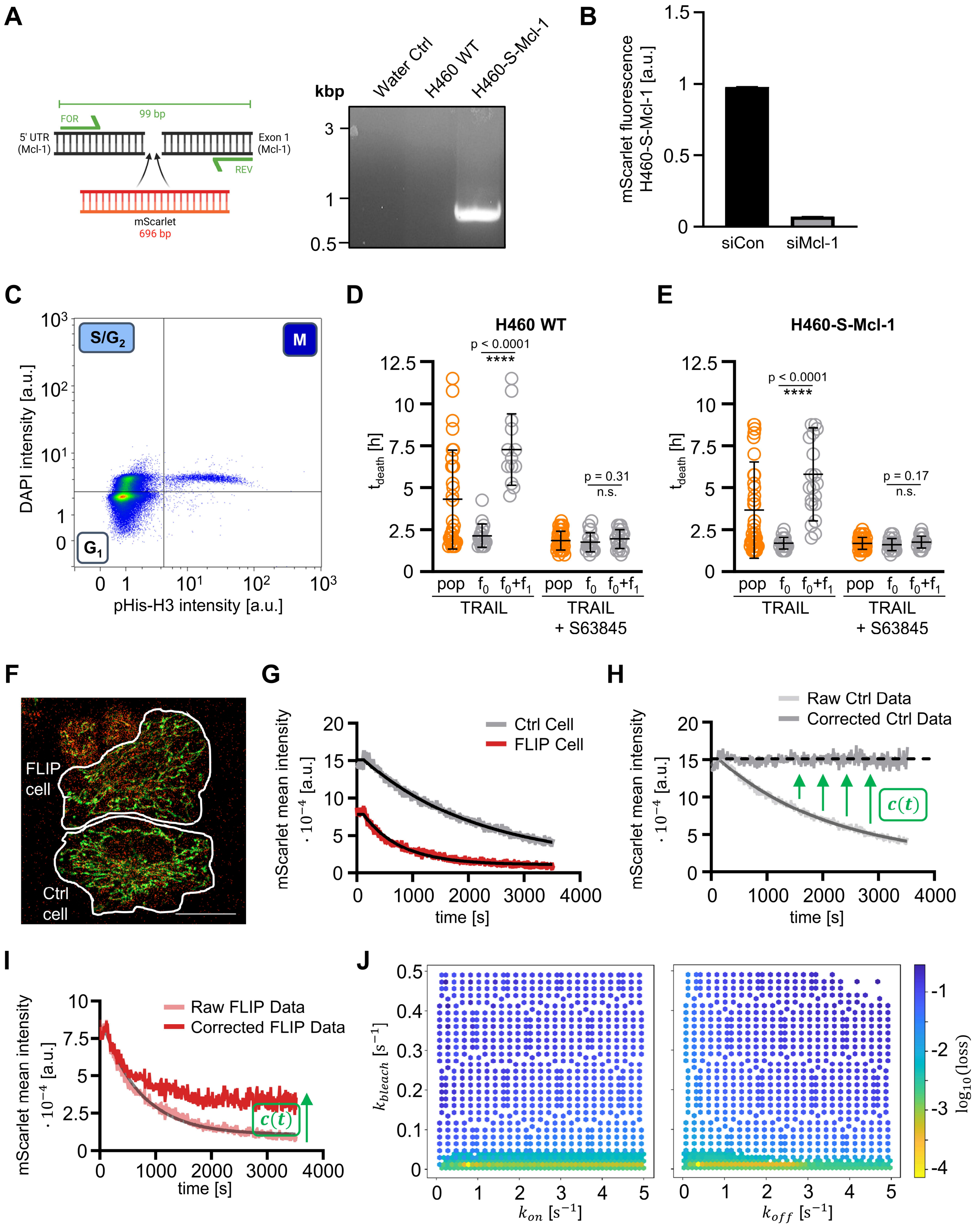


**Figure S2:**

A) Knock-in of the mScarlet-insertion into the Mcl-1 gene, and associated PCR product from genomic DNA amplification. The PCR product with mScarlet insertion is in the range of the expected size of 795 bp.

B) H460-S-Mcl-1 cells were treated either with unspecific control siRNA (*siCon*) or Mcl-1 specific siRNA (*siMcl-1*). Mean mScarlet fluorescence was measured 24 h after transfection via flow cytometry. n > 10 000 cells per group, one representative experiment from two independent repeats is shown.

C) H460-S-Mcl-1 cells were fixed, permeabilized, stained for DNA and pHis-H3, and analyzed by flow cytometry to separate cell cycle stages. One representative experiment from two independent repeats is shown.

D) H460 WT cells were either treated with 10 ng/ml Fc-scTRAIL (*TRAIL*) or co-treated with 10 ng/ml Fc-scTRAIL and 10 µM S63845 (*TRAIL + S63845*). After the treatment, the time until death ($t_{death}$) was analyzed for single cells. The whole population (*pop*) of tracked cells was separated into cells that did not undergo mitosis before dying (*f0*) and cells that went through mitosis before dying (*f0+f1*). n = 20 - 30 cells per group. Error bars represent mean ± standard deviation. *p*-values from an unpaired t-test with Welch's correction are depicted.

E) H460-S-Mcl-1 cells were treated and analyzed according to (D). n = 15 - 30 cells per group. Error bars represent mean ± standard deviation. *p*-values from an unpaired t-test with Welch's correction are depicted.

F) Experimental setup of the FLIP experiment with the FLIP cell that was bleached and the control cell as bleaching reference.

G) Decay of mScarlet-Mcl-1 intensity in the control cell (grey) or FLIP cell (red).

H) Graph depicting the function *c(t)* that was used to correct for unspecific photobleaching based on the loss in the control cell.

I) Graph depicting the application of *c(t)* to the measured decay of the FLIP cell.

J) Hexbinplot that shows the loss value (difference of the model to the experimental data) depending on the respective combination of $k_{on}/k_{off}$ and $k_{bleac}$ in the FLIP model.

**Figure S3**

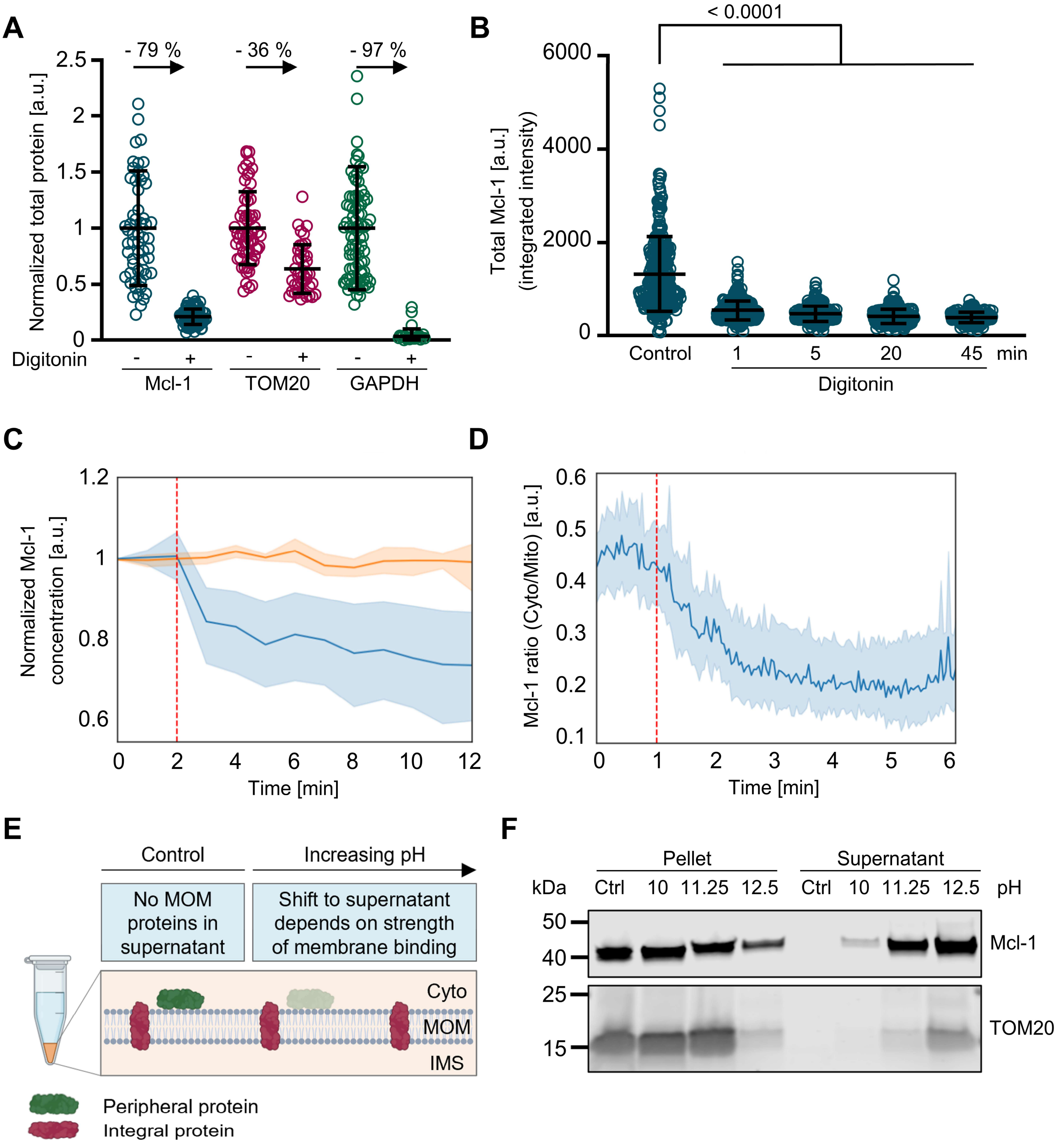


**Figure S3:**

A) Quantification of Mcl-1, TOM20 and GAPDH intensity in NCI-H460/geminin cells before (control) and after treatment with digitonin. Values were normalized to the control mean of the respective protein. Percent reductions are indicated. Data are from n = 34 - 82 cells per group,

B) Quantification of total Mcl-1 amounts in NCI-H460/geminin cells before (control) and after treatment with digitonin, fixation at various timepoints after treatment.

Error bars represent mean ± standard deviation. *p*-values from a One-Way ANOVA with Dunnett's T3 multiple comparisons post hoc test are depicted.

C) Quantification of Scarlet-Mcl-1 mean intensity (concentration) over time. The blue line represents the mean Mcl-1 signal in digitonin-treated cells (n = 16 cells from three independent experiments). The red dashed line indicates the time of digitonin addition. The orange line shows the mean Mcl-1 signal in untreated control cells imaged under identical conditions to account for photobleaching (n = 5 cells from three independent experiments). Fluorescence intensities were normalized to timepoint 0 and curve corrected to the bleaching control. Shaded areas indicate standard deviation across cells.

D) Quantification of the ratio between cytoplasmic and mitochondrial Mcl-1 concentrations in NCI H460/mScarlet cells (n = 16 cells) via super resolution live cell imaging. Shaded areas indicate standard deviation across cells. The red dashed line indicates the time of digitonin addition.

E) Schematic representation of the sodium carbonate extraction used to assess the strength of protein membrane association. Mitochondria were isolated and incubated in 100 mM $Na_2CO_3$ with different pH values. After an ultracentrifugation step, proteins that are loosely associated with membranes migrate into the supernatant as the pH increases, while tightly bound or integral proteins remain in the pellet.

F) Immunoblot analysis of Mcl-1 and TOM20 membrane association during sodium carbonate extraction. Purified mitochondria from NCI-H460/geminin cells were treated with $Na_2CO_3$ at the indicated pH and then centrifuged to collect the supernatant and the pellet fractions.

**Figure S4**

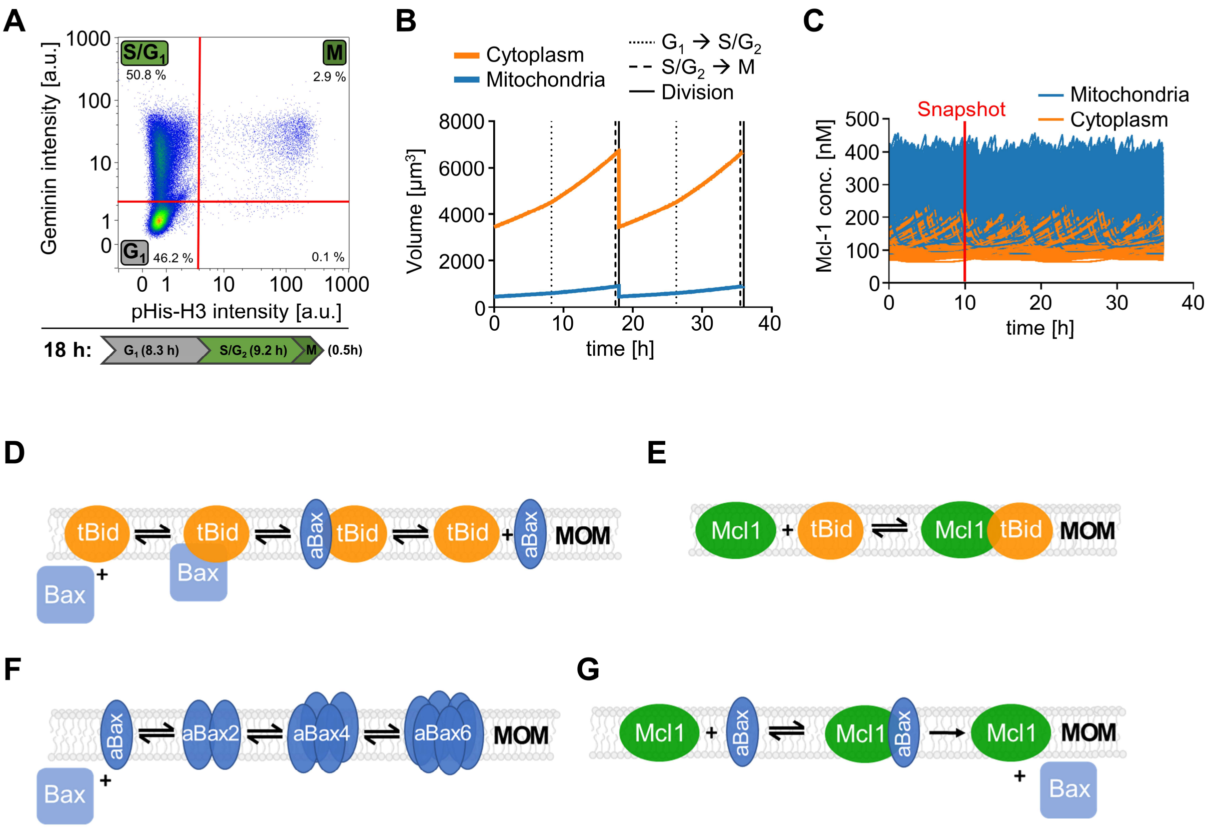


**Figure S4:**

A) NCI-H460/geminin cells were fixed, permeabilized and stained for pHis-H3, to separate cell cycle stages as indicated. The relative proportion of cells in each phase was used to resolve the 18 h of total cell cycle duration into the respective cell cycle phases and durations.

B) Implementation of volume changes. With cell cycle progression, both the cytoplasmic and the mitochondrial compartment grow, followed by resetting to starting values upon cell division.

C) A population of 2000 unsynchronized cells, their basal Mcl-1 expression and distribution ratio was simulated. At 10 h of the simulation, a population snapshot was taken for comparison to experimental data (see Figure 4D,E).

D) Scheme depicting the interactions between t-Bid and Bax/Bak in the apoptosis susceptibility model

E) Scheme depicting the interactions between t-Bid and Mcl-1 in the apoptosis susceptibility model

F) Scheme depicting the oligomerization of Bax/Bak in the apoptosis susceptibility model

G) Scheme depicting the interactions between Bax/Bak and Mcl-1 in the apoptosis susceptibility model

### Figure S5

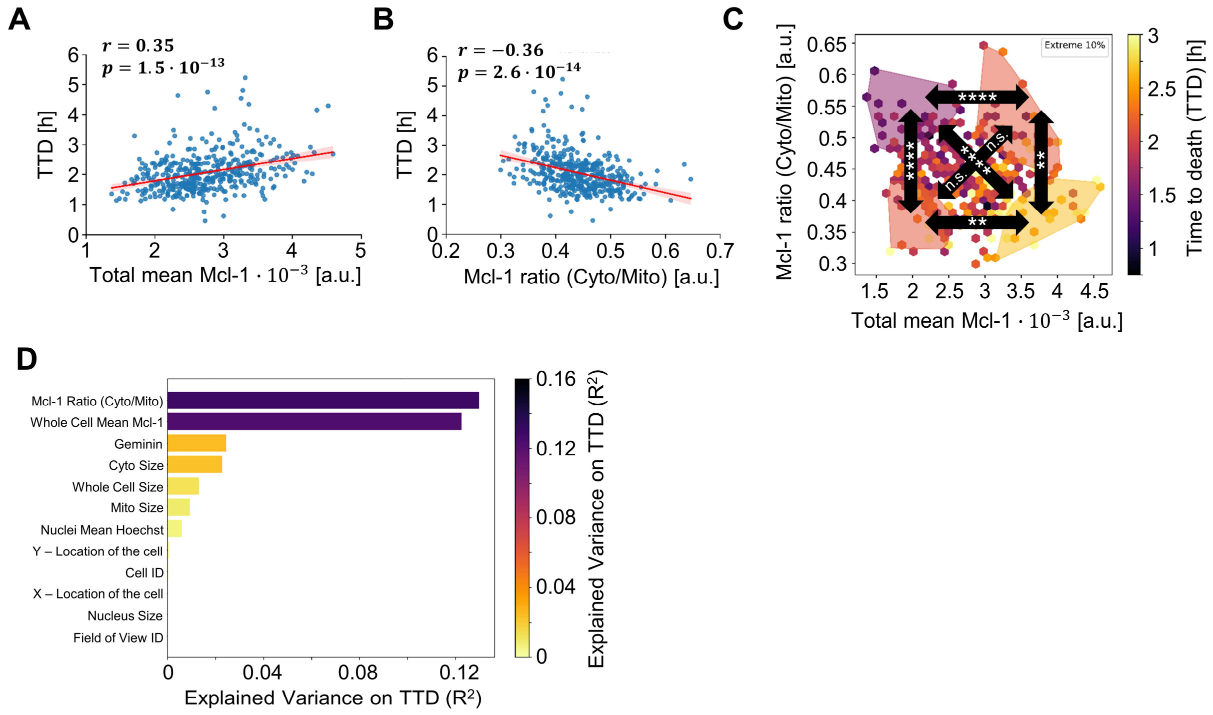


**Figure S5:**

A) Correlation between time to death (TTD) and total Mcl-1 expression, with the respective Pearson’s r and *p* value indicated.

B) Correlation between time to death (TTD) and Mcl-1 ratio, with the respective Pearson’s r and *p* value indicated.

C) All groups were compared using One Way ANOVA with Dunnett’s T3 multiple comparisons.

D) The variance ($R^2$) in the measured TTD that is explained through each of the experimentally available parameters is depicted.

**Figure S6**

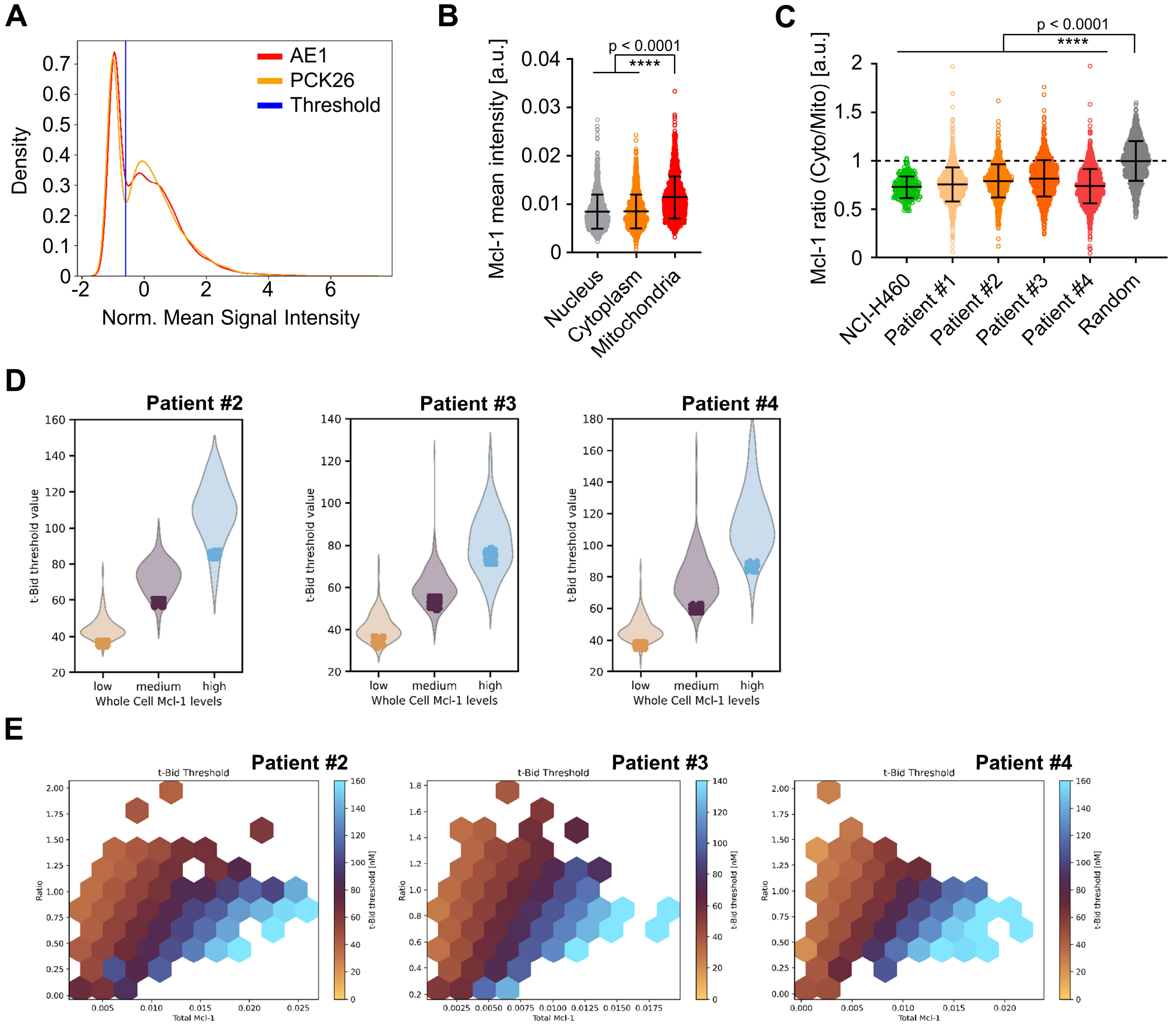


**Figure S6:**

A) Distribution of tumour marker intensities and the threshold value depicting the discrimination between tumour and non-tumour cells.

B) Subcellular concentrations of Mcl-1 in nuclei, cytoplasm and mitochondria of human tumour cells. Error bars represent mean ± standard deviation. *p*-value from a One-Way ANOVA with Dunnett's T3 multiple comparisons post hoc test are depicted.

C) Comparison of Mcl-1 cytosolic to mitochondrial distribution ratios between NCI-H460 cells, patient tumour samples, and a control of randomly assigned segmentation regions (The control represents randomly distributed mitochondrial regions within the cell). Error bars represent mean ± standard deviation. *p*-value from a One-Way ANOVA with Dunnett's T3 multiple comparisons post hoc test are depicted.

D) The t-Bid threshold values where 10% of Bax/Bak end up in pores were compared. Thresholds determined from only Mcl-1 expression amounts as input are depicted as solid filled dots. Thresholds derived from both Mcl-1 expression and distribution as inputs are depicted as violin plots.

E) Hexbinplot showing the simulated t-Bid thresholds of single tumour cells dependent on the Mcl-1 distribution ratio and the total Mcl-1 expression. Thresholds increase along both axes.
